# Spatial mapping of α-synuclein seeding activity and pathology using immunofluorescence-integrated quiescent seed amplification

**DOI:** 10.1101/2025.03.26.645411

**Authors:** Hengxu Mao, Yaoyun Kuang, Zerui Wang, Jin Li, Wei Dai, Hao Lin, Tingting Gan, Zhengxiao He, Wen-quan Zou, Chao Ma, Yue Huang, Jinkun Huang, Ping-Yi Xu

## Abstract

Seed amplification assays sensitively detect misfolded α-synuclein (αSyn) templating activity in homogenates but erase tissue coordinates, whereas Ser129-phosphorylated αSyn (pS129) immunostaining preserves anatomy without measuring functional seeding competence. Here we report multiplexed immunofluorescence-integrated quiescent seed amplification (IF-QSAA), coupling in situ QSAA with multichannel immunolabelling on the same brain section. Combining mouse αSyn monomer, high sulfate and 70 °C quiescent incubation under an agarose overlay generates localized, fibrillar thioflavin T-positive products with high seed specificity. Across preformed-fibril-inoculated mouse brains and human Parkinson’s disease (PD) tissue, QSAA reveals white-matter seeding enrichment, resolves hemispheric asymmetry earlier than pS129, and recovers endogenous seeding signals from archival formalin-fixed PD tissue lacking pS129 immunoreactivity. Using a QSAA-first workflow with 80 °C citrate retrieval and spectral unmixing, IF-QSAA identifies selective neuronal and microglial associations and resolves core–halo QSAA–pS129 dual-positive pairs. IF-QSAA thus co-maps functional seeding and molecular histopathology.

**Teaser:** Same-section seed amplification and immunostaining map functional α-synuclein seeds across cellular neighbourhoods and single inclusions.

## Introduction

The misfolding, aggregation and templated propagation of α-synuclein (αSyn) are defining molecular hallmarks of Parkinson’s disease (PD), dementia with Lewy bodies and multiple system atrophy [1,2]. Pathological αSyn assemblies act as conformational templates that recruit soluble monomer into growing fibrils, a functional property termed seeding activity [3–6]. Comparing, within tissue, the amplifiable seeding activity of αSyn, its molecular pathological state and its cellular microenvironment requires an assay that captures these together while preserving their spatial coordinates. However, current methodologies force a trade-off between functional sensitivity and spatial resolution.

On one side of this methodological divide, histological immunostaining, most commonly targeting αSyn phosphorylated at serine 129 (pS129), preserves tissue architecture and resolves the regional distribution, cellular neighbourhood and morphology of inclusions [7,8]. Yet immunoreactivity measures the presence and steric accessibility of a post-translational modification on deposited material, not whether that material retains catalytic competence to template new fibril growth [9–12]. Experimental evidence indicating that Ser129 phosphorylation can occur after initial aggregation and may even inhibit further fibril elongation underscores that histological deposition and seeding activity are non-interchangeable dimensions of pathology [10,11]. On the other side, seed amplification assays (SAAs), including real-time quaking-induced conversion (RT-QuIC) and protein misfolding cyclic amplification (PMCA), directly measure templating activity with femtogram-to-attogram sensitivity [13–16]. Standard SAAs rely on cyclic mechanical shaking or sonication to fragment growing fibrils and multiply active fibril ends. Because this mechanical regime requires tissue homogenization or fluid extraction, a positive bulk readout cannot be mapped back to a specific tissue region or cellular structure.

Efforts to bridge this spatial–functional gap have recently begun. We previously introduced the quiescent seed amplification assay (QSAA) and showed that amplification-derived thioflavin T (ThT) fluorescence can be generated without mechanical fragmentation on intact tissue sections [17], and an independent in situ seeding immunodetection strategy recently used a tagged recombinant substrate to visualize newly recruited αSyn in human tissue [18]. Despite these advances, turning QSAA based on direct ThT readout into a quantitative spatial analysis method that can be integrated on the same section with multichannel immunofluorescence (IF) still requires establishing the corresponding reaction workflow and analytical methods experimentally. On the one hand, seed-dependent amplification without mechanical fragmentation and the spatial retention of amplified products within tissue need to be evaluated. On the other hand, standalone QSAA is a single-channel assay, and comparing it with pS129 or cellular markers on adjacent sections makes it difficult to achieve one-to-one correspondence at the level of individual cells or micrometre-scale deposits. It is therefore necessary to evaluate the compatibility of QSAA and immunofluorescence on the same section and to examine how assay order, antigen retrieval and imaging channels affect both types of signal.

To establish this same-section combined detection method, we developed a multiplexed same-section IF-QSAA quantitative spatial analysis framework spanning reaction kinetic engineering, anti-convective in situ tissue design, multi-channel protocol optimization and spatial statistical null models. We first define the kinetic basis of a highly specific fragmentation-free amplification window and establish an atlas-registered workflow for in situ QSAA. We then perform whole-brain grey- and white-matter comparisons and early post-inoculation time-course analyses in an exogenous mPFF-inoculated mouse model, and validate the amplification of endogenous seeds in human PD brain tissue. Next, we quantify the effects of assay order, antigen retrieval and spectral bleed-through to establish a standardized same-section IF-QSAA protocol. Finally, we combine this method with spatial permutation and translation null models to achieve co-registered quantitative analysis of seeding activity alongside cellular markers and pS129 pathology at the scales of cellular microenvironments and individual pathological objects.

## Results

### 1. Establishing a quiescent amplification reaction system without mechanical fragmentation

To amplify tissue-associated seeds in situ without destroying section integrity, a reaction must achieve rapid fibril elongation without cyclic mechanical fragmentation while suppressing spontaneous (unseeded) nucleation over the assay duration (Fig. 1a). In conventional SAAs, periodic sonication or shaking repeatedly breaks fibrils at a local elongation plateau, multiplying growth-competent ends [19,20] (Fig. 1a,b). A quiescent system instead relies primarily on seed-end elongation and secondary nucleation, requiring biochemical reconstruction of its kinetic driving forces [17] (Fig. 1b).

**Fig. 1.**
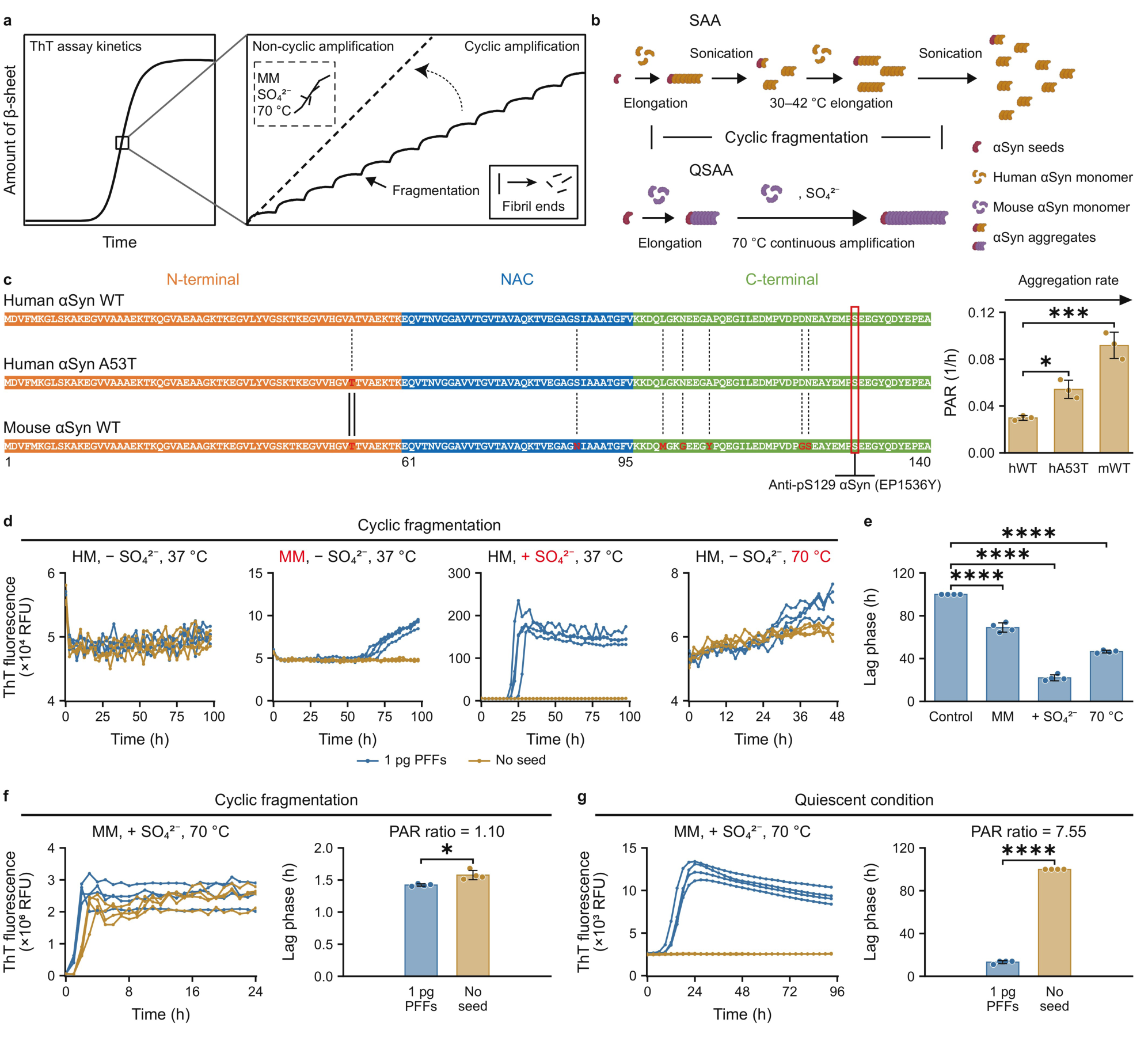
Mouse monomer, sulfate and 70 °C support seeded amplification without fragmentation. **a**, Scheme: each step of the cyclic thioflavin T (ThT) curve follows fragmentation creating fibril ends; dashed line, non-cyclic amplification, boxed conditions. **b**, Seed amplification assay (SAA): sonication fragments a fibril grown from an α-synuclein (αSyn) seed and human monomer (HM). Quiescent SAA (QSAA): continuous growth with mouse monomer (MM), sulfate (SO_4_^2-^) and 70 °C. **c**, Left, αSyn alignment: human wild-type (hWT), A53T (hA53T) and mouse wild-type (mWT); red, residues differing from hWT and the Ser129 box, epitope of anti-pS129 antibody EP1536Y (αSyn phosphorylated at Ser129); NAC, non-amyloid-β component. Right, protein aggregation rate (PAR) per monomer. **d**, ThT fluorescence under cyclic fragmentation; conditions above each plot, factor changed from the reference (left) in red; each line, one well; blue, 1 pg preformed fibrils (PFFs); ochre, no seed. **e**, Lag phase of seeded wells in **d**. **f**, **g**, MM, +SO_4_^2-^ and 70 °C, cyclic fragmentation (**f**) or quiescent (**g**): ThT fluorescence and lag phase, seeded and unseeded wells; PAR ratio, seeded/unseeded. Bars, mean ± s.d.; circles, wells; n = 3 (**c**) or 4 (**e**–**g**) wells; one-way ANOVA with Dunnett’s test against hWT (**c**) or Control (**e**); two-tailed unpaired *t*-test (**g**); *P < 0.05, ***P < 0.001, ****P < 0.0001.

We purified full-length recombinant αSyn monomer substrates by anion-exchange chromatography; the main SDS–PAGE band ran at 17.3 kDa with ≥ 95.2% purity (representative lane) and a mean of 93.6 ± 1.6% across three fractions (Supplementary Fig. 1a,b). Human wild-type (hWT) and mouse wild-type (mWT) αSyn differ by seven amino-acid substitutions across the N-terminal, NAC and C-terminal domains, while the Ser129 phosphorylation site recognized by EP1536Y is conserved (Fig. 1c, left). In unseeded reactions under cyclic shaking, the protein aggregation rate (PAR, the reciprocal of the lag phase) was highest for mWT (0.092 h^-1^), then the disease-associated human A53T variant (hA53T, 0.054 h^-1^) and lowest for hWT (0.030 h^-1^) [14,21,22] (Fig. 1c, right). Preformed fibril (PFF) seeds sonicated from human (hPFF) and mouse (mPFF) fibrils showed short fibrillar morphology by negative-stain transmission electron microscopy (TEM), mPFFs with a shorter, narrower length distribution (median 45 nm) than hPFFs (median 76 nm; Supplementary Fig. 1c–e).

We next dissected how monomer species [14,21,22], kosmotropic sulfate ions (SO_4_^2-^) [23–26] and elevated temperature (70 °C) [27,28] individually affect amplification kinetics when seeded with 1 pg PFFs under cyclic fragmentation (Fig. 1d). Whereas the reference condition (hWT monomer, no sulfate, 37 °C) failed to distinguish seeded from unseeded wells within 100 h, mWT monomer (lag phase ∼64–74 h), added sulfate (∼20–26 h) or 70 °C incubation (∼45–48 h) each significantly accelerated the onset of seeded reactions (n = 4; Fig. 1d,e). However, with all three factors combined (mWT monomer + SO_4_^2-^+ 70 °C), cyclic fragmentation nucleated unseeded reactions almost as rapidly as seeded ones, collapsing the seeded-to-unseeded PAR ratio to 1.102 and abolishing analytical specificity (Fig. 1f).

Removing mechanical fragmentation from the same high-reactivity formulation restored a wide, seed-specific operating window (Fig. 1g). Quiescent reactions seeded with 1 pg mPFFs amplified after ∼11–14 h, whereas all unseeded control wells remained at baseline throughout the 93–96 h run, increasing the seeded-to-unseeded PAR ratio to 7.547 (P < 0.0001, n = 4; Fig. 1g). TEM of quiescent endpoints confirmed that fluorescence increases corresponded to bona fide fibrillar assemblies (dense electron-opaque cores fringed by radiating fibrils and open fibril networks); unseeded control grids contained no fibrils (Supplementary Fig. 1f–i). Omitting fragmentation is thus a kinetic requirement that suppresses spontaneous nucleation in the high-elongation-rate system combining mouse monomer, high sulfate and 70 °C.

### 2. Application of the quiescent amplification reaction system to brain sections from mPFF-inoculated mice

To translate solution-phase quiescent amplification to intact brain sections while preventing micro-convective displacement of products, we developed a hydrogel-confined slide workflow (Fig. 2a). Brain tissue fixed in 4% paraformaldehyde (PFA) was cryosectioned at 14 μm, post-fixed in 4% PFA and covered with a ∼1-mm-thick semi-solid 1% agarose layer [29], which confines newly formed fibrillar products while monomer and dye diffuse to the tissue surface. Each section received 300 μL of reaction mixture containing 1 mg mL^-1^ mWT αSyn monomer, approximately 800 mM Na_2_SO_4_ and 40 μM ThT. Slides were incubated quiescently in a humidified chamber at 70 °C for 8 h and imaged directly (Fig. 2a).

**Fig. 2.**
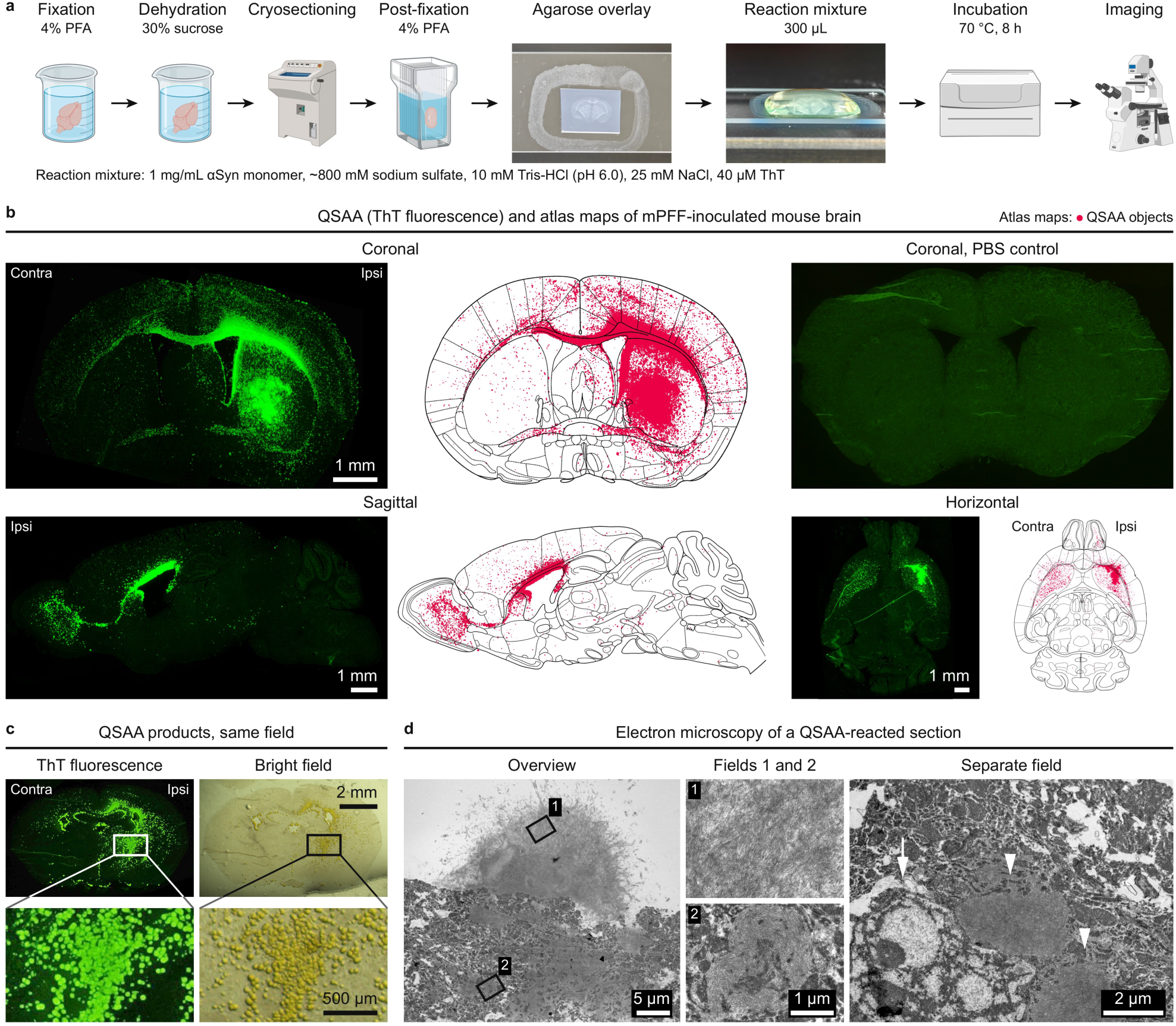
QSAA on brain sections yields fibrillar, thioflavin T-positive products that map the anatomy of the inoculated mouse brain. **a**, Workflow of the QSAA: brains are fixed and cryosectioned; sections are post-fixed in 4% paraformaldehyde (PFA), overlaid with agarose and reaction mixture, incubated at 70 °C for 8 h and imaged. The mixture contains αSyn monomer and ThT, composition printed beneath the workflow. Photographs, the agarose-overlaid section and a mixture-covered slide. **b**, QSAA on coronal, sagittal and horizontal mouse brain sections 180 days after unilateral striatal inoculation of mouse αSyn preformed fibrils (mPFF). Green, ThT fluorescence. Atlas maps beside each micrograph show its QSAA objects as crimson dots. Hemispheres are labelled contralateral (Contra) and ipsilateral (Ipsi) to the inoculation; the sagittal section is ipsilateral. Rostral, left in the sagittal and up in the horizontal section. Control, a coronal section from a phosphate-buffered saline (PBS)-inoculated mouse. Sagittal and horizontal series, Supplementary Fig. 2; coronal levels analysed in detail, Fig. 3 and Supplementary Fig. 3. **c**, ThT fluorescence and bright-field images of one field of a coronal section after QSAA; boxes mark regions enlarged below. **d**, Transmission electron micrographs of a QSAA-reacted section: outlines 1 and 2 mark fields 1 and 2; arrowheads in the separate field mark fibrillar aggregates within the tissue. Scale bars, 1 mm (**b**), 2 mm (**c**, upper), 500 μm (**c**, lower), 5 μm (**d**, overview), 1 μm (**d**, fields 1 and 2) and 2 μm (**d**, separate field).

We tested whether this configuration preserves three-dimensional neuroanatomical organization in mice 180 days post-inoculation (dpi) after unilateral striatal injection of mPFFs [5,30]. Across coronal, sagittal and horizontal planes, in situ QSAA generated bright, anatomically structured ThT-positive signals segmented into individual objects and registered to mouse brain atlas coordinates (Fig. 2b). Signal was densest around the ipsilateral striatal injection site and extended along connected grey-matter territories and interhemispheric commissures (the corpus callosum and anterior commissure) into the contralateral hemisphere, with sagittal views clearly showing dissemination along the olfactory limb of the anterior commissure toward the olfactory bulb. Sections from phosphate-buffered saline (PBS)-inoculated control mice showed no specific particulate signal (Fig. 2b). Complete serial mapping across 16 sagittal sections (lateral to medial; Supplementary Fig. 2a) and 12 horizontal sections (dorsal to ventral; Supplementary Fig. 2b) confirmed that the ipsilateral predominance, cortical dispersion and tract-aligned continuity were reproducible across orthogonal axes.

To confirm that the ThT signal marks physical amplification products, we performed correlative bright-field and ultrastructural imaging. Bright-field microscopy of QSAA-reacted sections revealed discrete, yellow-brown granular deposits whose spatial coordinates and morphologies matched the ThT-positive puncta in the same field (Fig. 2c). In situ TEM of ultrathin sections from QSAA-reacted brain slices demonstrated electron-dense product masses with radiating amyloid-like fibrils and bundles of fibrillar aggregates embedded directly within the tissue matrix (Fig. 2d).

### 3. Whole-brain atlas registration and regional benchmarking of QSAA against pS129

To evaluate what spatial information QSAA adds to standard immunohistopathology, we benchmarked whole-section QSAA object maps against pS129 IF on anatomically matched coronal sections at 180 dpi (Fig. 3a; Supplementary Fig. 3a–e). At the Bregma +1.46 mm striatal injection level, a five-step registration and segmentation pipeline yielded 1,234 QSAA-positive objects (teal dots) and 1,513 pS129-positive objects (magenta dots; Fig. 3a). We mapped 4,459 QSAA objects and 3,400 pS129 objects across five rostro-caudal levels (Bregma +1.46, +1.11, -0.92, -1.51 and - 3.07 mm; Supplementary Fig. 3a–e).

**Fig. 3.**
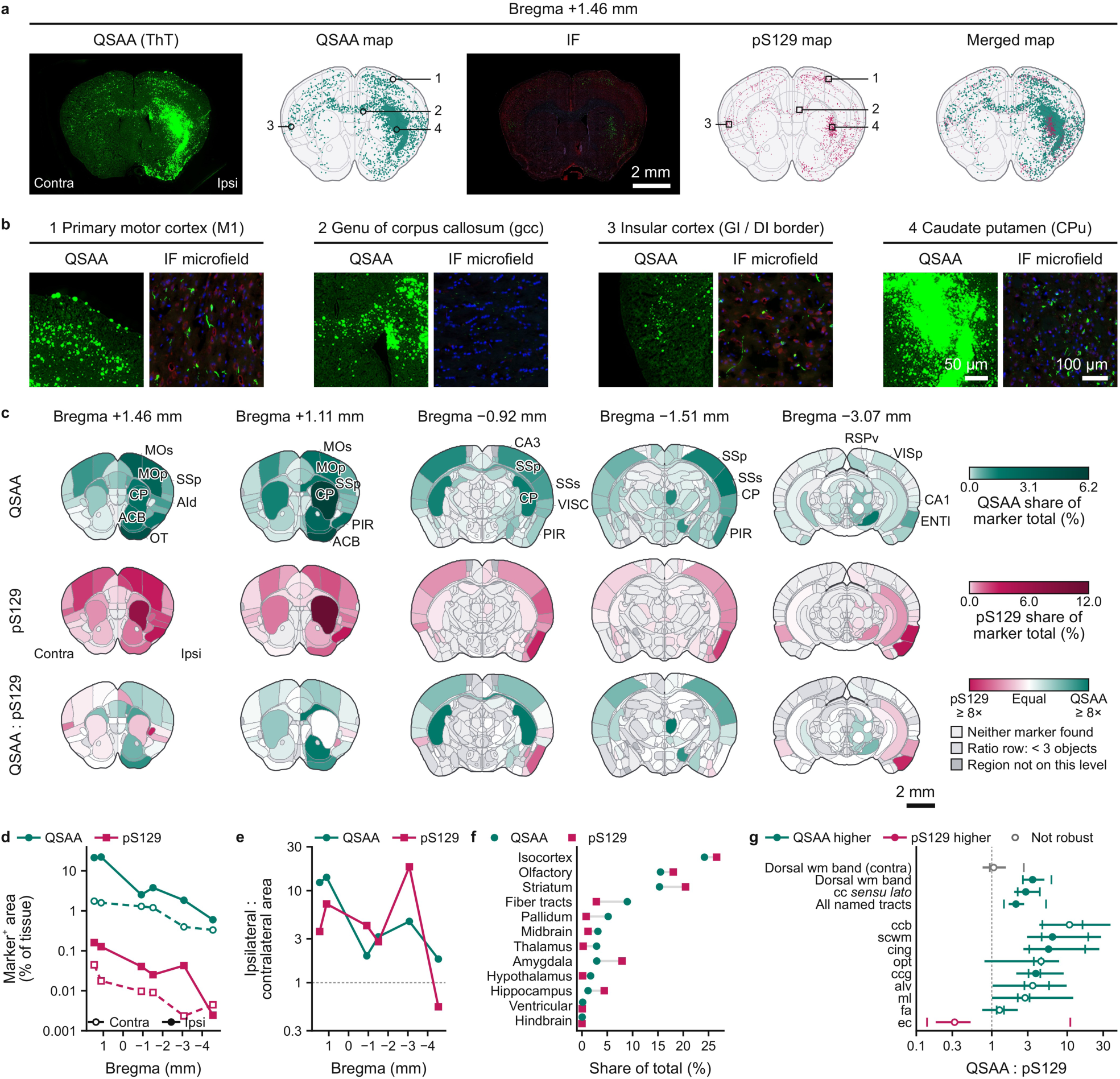
QSAA products and pS129 immunofluorescence differ in regional distribution across the mPFF-inoculated mouse brain. **a**, One coronal level (Bregma +1.46 mm) of mouse brain 180 days after inoculation of mouse αSyn mPFF into one striatum, in five analysis steps, left to right: QSAA, whole-section QSAA products, ThT fluorescence (green); QSAA map, those objects on the atlas map, one teal dot each; IF, the matched section stained for αSyn phosphorylated at serine 129 (pS129, green), neuronal nuclei antigen (NeuN, red) and 4′,6-diamidino-2-phenylindole (DAPI, blue); pS129 map, those objects (magenta dots); merged map, both together. Rings (QSAA map) and squares (pS129 map) numbered 1–4, the four fields of **b** in the section outline. Contra and Ipsi hemispheres, left and right throughout. **b**, The four fields of **a**, by region: QSAA, enlarged ThT fluorescence; IF microfield, the same region on the matched IF section, channels as in **a**. Region and tract abbreviations, Supplementary Table 2. Other five levels and their fields, Supplementary Fig. 3. **c**, Regional distribution of both markers at five coronal levels, rostral to caudal; Bregma coordinates, Allen reference atlas. QSAA and pS129 rows, each region per hemisphere coloured by its percentage of that marker’s total object area. QSAA : pS129 row, the share ratio (teal, QSAA higher; magenta, pS129 higher; white, equal; scale ends at 8×). Grey fills, as in the key. **d**–**g**, Quantification; teal, QSAA; magenta, pS129. **d**, Marker-positive area as a percentage of tissue area per hemisphere at the six coronal levels (**a** and Supplementary Fig. 3). Circles, QSAA; squares, pS129. Filled symbols and solid lines, ipsilateral; open symbols and dashed lines, contralateral. **e**, Ipsilateral-to-contralateral ratio of the areas in **d**; dotted line, 1. **f**, Share of each marker’s total object area in twelve ipsilateral territories, pooled over the levels of **c**; grey bars join the markers. **g**, QSAA : pS129 enrichment in white-matter tracts, a tract’s QSAA share divided by its pS129 share. Circles, ratio; bars, 95% bootstrap confidence interval (CI); ticks, the leave-one-level-out range; dotted line, 1. Filled circles, CI and leave-one-level-out range exclude 1; open circles, not. Teal, ratio above 1; magenta, below 1; grey, contralateral hemisphere; other rows, ipsilateral. Scale bars, 2 mm (**a**, IF section; **c**), 50 μm (**b**, QSAA) and 100 μm (**b**, IF microfields).

Paired regional microfields showed that both assays detected pathology in grey-matter structures including the primary motor cortex (M1), granular/dysgranular insular cortex (GI/DI) and caudate putamen (CPu), but diverged sharply in white matter: the genu of the corpus callosum (gcc), internal capsule (ic) and nigrostriatal bundle (ns) contained dense QSAA amplification products where matched pS129 IF showed only sparse or isolated neuritic profiles (Fig. 3b; Supplementary Fig. 3a–e). Regional share maps across the five coronal levels confirmed this brain-wide (Fig. 3c). When normalized to each marker’s own total object area and summarized across 12 anatomical divisions, both readouts were similarly distributed across grey-matter divisions, with the ipsilateral isocortex holding 24% of QSAA and 27% of pS129 object area and the ipsilateral striatum 15% and 20%, respectively (Fig. 3f), and both exhibited a consistent rostral-to-caudal gradient and ipsilateral predominance (ipsilateral-to-contralateral area ratio of 2.0–14.0 for QSAA and 2.8–18.2 for pS129 across the five rostral levels; Fig. 3d,e). In absolute terms, QSAA-positive area exceeded pS129-positive area by one to two orders of magnitude at every level (Fig. 3d).

Within the fibre-tract division, relative composition differed between the assays (Fig. 3f,g): a tract-by-tract bootstrap analysis of the QSAA-to-pS129 object-share ratio demonstrated a 3.5-fold enrichment across the pooled ipsilateral dorsal white-matter band comprising the corpus callosum body (ccb), genu (ccg), anterior forceps (fa), supra-callosal white matter (scwm) and cingulum bundle (cing) (95% bootstrap confidence interval [CI] 2.5–5.0), driven by 3.8-to 6.4-fold enrichments in ccg, cing and scwm (Fig. 3g). By contrast, the ratio for the external capsule (ec) fell below 1, and in the contralateral dorsal white-matter band the within-hemisphere relative share ratio was close to 1 (ratio 1.06, 95% CI 0.76–1.54; Fig. 3g). Spatial QSAA thus captures tract-associated seeding activity that conventional pS129 immunostaining under-represents.

### 4. Early post-inoculation time course in mice and endogenous seed detection in human PD brain

We next tested whether spatial QSAA can still detect seeding activity where conventional staining is faint or its epitope degraded: early post-inoculation stages in mice and post-mortem human brain tissue. At 30 dpi in mPFF-inoculated mice, benchmarking against pS129 IF and thioflavin S (ThS) amyloid staining showed that QSAA labelled a substantially wider territory across the bilateral cortex, corpus callosum and striatum than either comparator (Fig. 4a). In the ipsilateral cortex, QSAA-positive area reached 69.0%, significantly exceeding both pS129 IF (30.8%, adjusted P = 1.3 × 10^-6^) and ThS (22.8%, adjusted P = 2.5 × 10^-7^, n = 4; Fig. 4b). Because QSAA recruits exogenous monomer to generate new ThT-positive products whereas IF and ThS only label pre-existing deposits, this area difference reflects the detection and spatial amplification of seed signals.

**Fig. 4.**
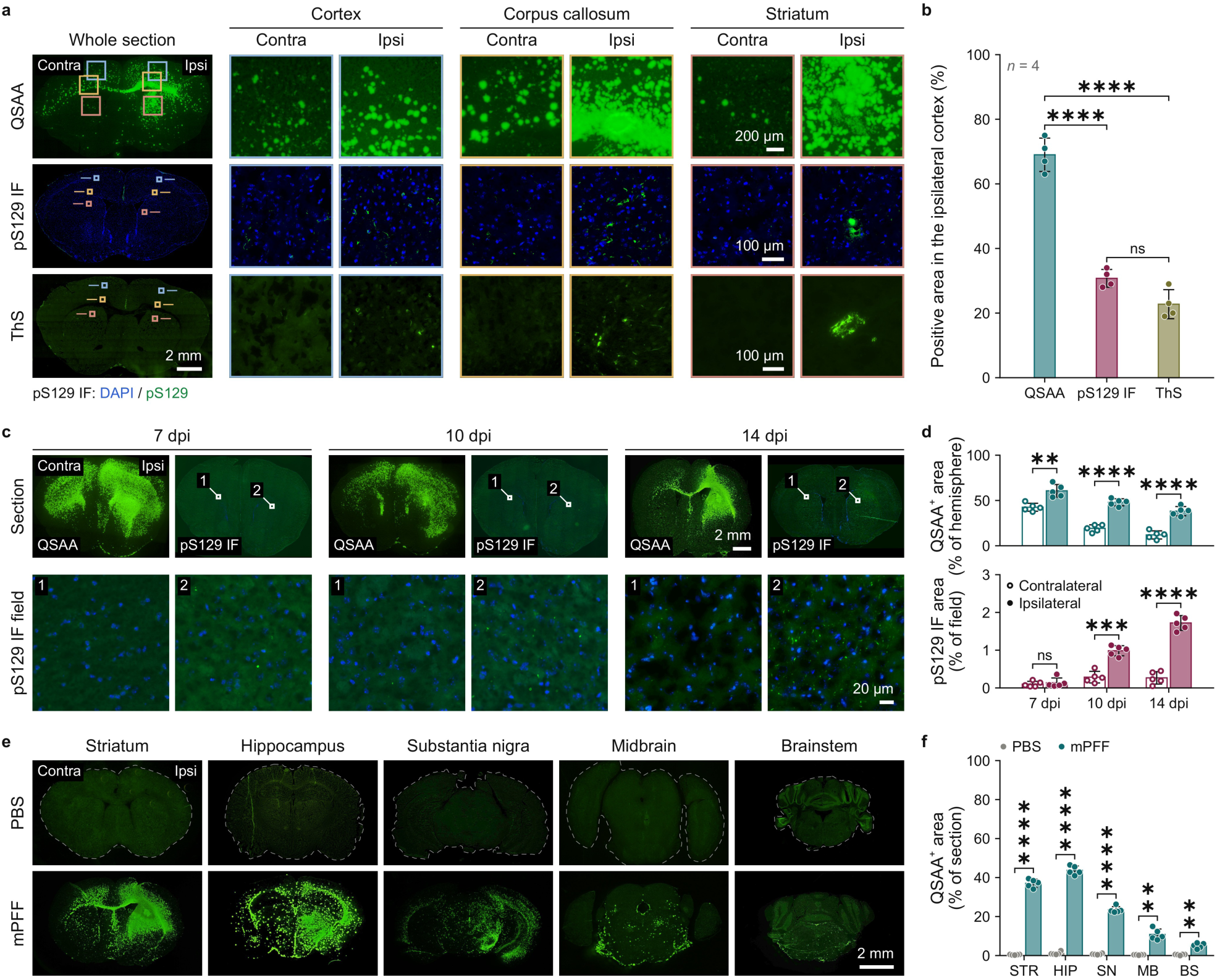
QSAA marks a larger area than pS129 immunofluorescence or thioflavin S and is positive from 7 days after inoculation. **a**, Coronal mouse brain sections 30 days post-inoculation (dpi) with mouse αSyn mPFF, processed for QSAA (green), IF for pS129 (green; DAPI, blue) or thioflavin S (ThS; green). Coloured boxes mark cortical, callosal and striatal fields at right in the same colour, Contra and Ipsi to the inoculation. **b**, Positive area, ipsilateral cortex, 30 dpi. Bars, means; dots, individual values; error bars, s.d.; n = 4 per method; one-way ANOVA with Tukey’s multiple-comparisons test. **c**, Coronal sections, QSAA or pS129 IF, 7, 10 and 14 dpi. Numbered boxes on the pS129 IF sections, fields below (1, Contra; 2, Ipsi). **d**, QSAA-positive area per hemisphere (top) and pS129-positive area per field (bottom), 7, 10 and 14 dpi. Bars, means; dots, individual values (open, Contra; filled, Ipsi); error bars, s.d.; n = 5 per group; two-tailed unpaired *t*-test with Welch’s correction, Contra versus Ipsi, Holm-adjusted over the three time points per plot. **e**, QSAA at 14 dpi at five rostro-caudal levels in PBS- or mPFF-inoculated mice. Dashed lines, PBS section outlines. **f**, QSAA-positive area per section at the five levels in **e**: striatum (STR), hippocampus (HIP), substantia nigra (SN), midbrain (MB) and brainstem (BS). Bars, means; dots, individual values (grey, PBS; teal, mPFF); error bars, s.d.; n = 5 per group; two-tailed unpaired *t*-test with Welch’s correction, PBS versus mPFF, Holm-adjusted over the five levels. *P < 0.05, **P < 0.01, ***P < 0.001, ****P < 0.0001; ns, not significant. Scale bars, 2 mm (whole sections, **a**, **e**; 14 dpi QSAA, **c**), 200 μm (QSAA fields, **a**), 100 μm (pS129 IF and ThS fields, **a**), 20 μm (14 dpi fields, **c**).

In an early time series at 7, 10 and 14 dpi, pS129 IF signal was extremely sparse and did not show a significant bilateral difference until 10–14 dpi, whereas QSAA resolved ipsilateral-versus-contralateral asymmetry at 7 dpi (P < 0.01, n = 5; Fig. 4c,d). Across five rostro-caudal levels at 14 dpi, mPFF-inoculated brains showed QSAA-positive area above the negligible signal of PBS-inoculated controls in striatum (STR), hippocampus (HIP), substantia nigra (SN), midbrain (MB) and brainstem (BS) (P < 0.01 to P < 0.0001, n = 5), falling from 43.7% of section area at the hippocampal level to 5.0% at the brainstem level (Fig. 4e,f). This steep gradient demonstrates that QSAA captures amplification-competent material before conventional pS129 histology forms overt deposits.

To validate the method on endogenous seeds formed during human disease, where exogenous inoculum is absent, we applied QSAA to frozen sections from patients with PD (n = 7), a formalin-fixed archival PD specimen (FF-PD, n = 4, technical replicates), a tauopathy specificity control (progressive supranuclear palsy, PSP, n = 6) and healthy controls (HC, n = 6) (Supplementary Fig. 4a,b). QSAA generated dense positive signal in both PD and FF-PD (both P < 0.0001 vs. HC), whereas PSP and HC remained at baseline (Supplementary Fig. 4a,b). In the >20-year FF-PD specimen, pS129 immunoreactivity was completely lost (positive area not significantly different from HC), whereas in situ QSAA after heat pretreatment recovered αSyn seeding activity (Supplementary Fig. 4a,b). Solution-phase RT-QuIC on homogenates from the same cases confirmed rapid amplification in PD and FF-PD (PAR ≈ 0.11–0.16 h^-1^) and flat baselines in HC and PSP over 16 h (n = 3; Supplementary Fig. 4c,d). Across five sampling sites, PD sections showed elevated QSAA signal in both striatal (29.5–32.2%, P < 0.0001) and both cortical (5.8–8.9%, P < 0.01 and P < 0.0001) regions compared with HC, but no signal above baseline in the human corpus callosum (not significant vs. HC, n = 5; Supplementary Fig. 4e,f), confirming the regional specificity of endogenous seed detection.

Finally, to test whether QSAA products nucleate and grow in place, we performed hourly time-lapse imaging (0–5 h) on human PD brain tissue during the reaction (Supplementary Fig. 4g,h). QSAA-positive area within regions of interest increased monotonically from 0.05% at 0 h to 12.7– 14% at 5 h (n = 5 regions; Supplementary Fig. 4g,h, left), while the drift-corrected centroid displacement of tracked objects remained at a median of 1.5–2.0 μm h^-1^ (less than a single image pixel, 2.78 μm; Supplementary Fig. 4h, right). Hydrogel-confined amplification products thus establish and expand locally within the tissue.

### 5. Workflow compatibility and signal calibration of same-section IF-QSAA

While standalone QSAA maps seeding activity at the regional scale, comparing seeding competence with cellular markers or pS129 within individual structures requires both assays on the same tissue section. Engineering evaluation revealed three mutually constraining physicochemical barriers to directly combining QSAA and IF in sequence.

Spectral measurements showed that ThT fluorescence (excited at 488 nm and read in the FITC window) overlaps completely with FITC; its emission tail extends into orange-red and far-red windows, and scattering components enter the nominal DAPI window (Fig. 5a). Single-source calibration on tissue showed QSAA products coupling into the DAPI channel at 49.57 ± 5.91%, the Texas Red channel at 23.94 ± 2.55% and the Cy5 channel at 5.91 ± 1.00% of their FITC-channel intensity, whereas DAPI, Alexa Fluor 555 and Alexa Fluor 647 showed no detectable cross-coupling (n = 4; Fig. 5b); bleed-through at amplification sites matched these ratios. We therefore reserved the 488-nm channel exclusively for ThT, assigned antibodies to the Alexa Fluor 555 (Texas Red) and Alexa Fluor 647 (Cy5) channels, and implemented pixel-wise linear unmixing, which eliminated product-shaped bleed-through while preserving nuclear and immunolabelled structures (Fig. 5c).

**Fig. 5.**
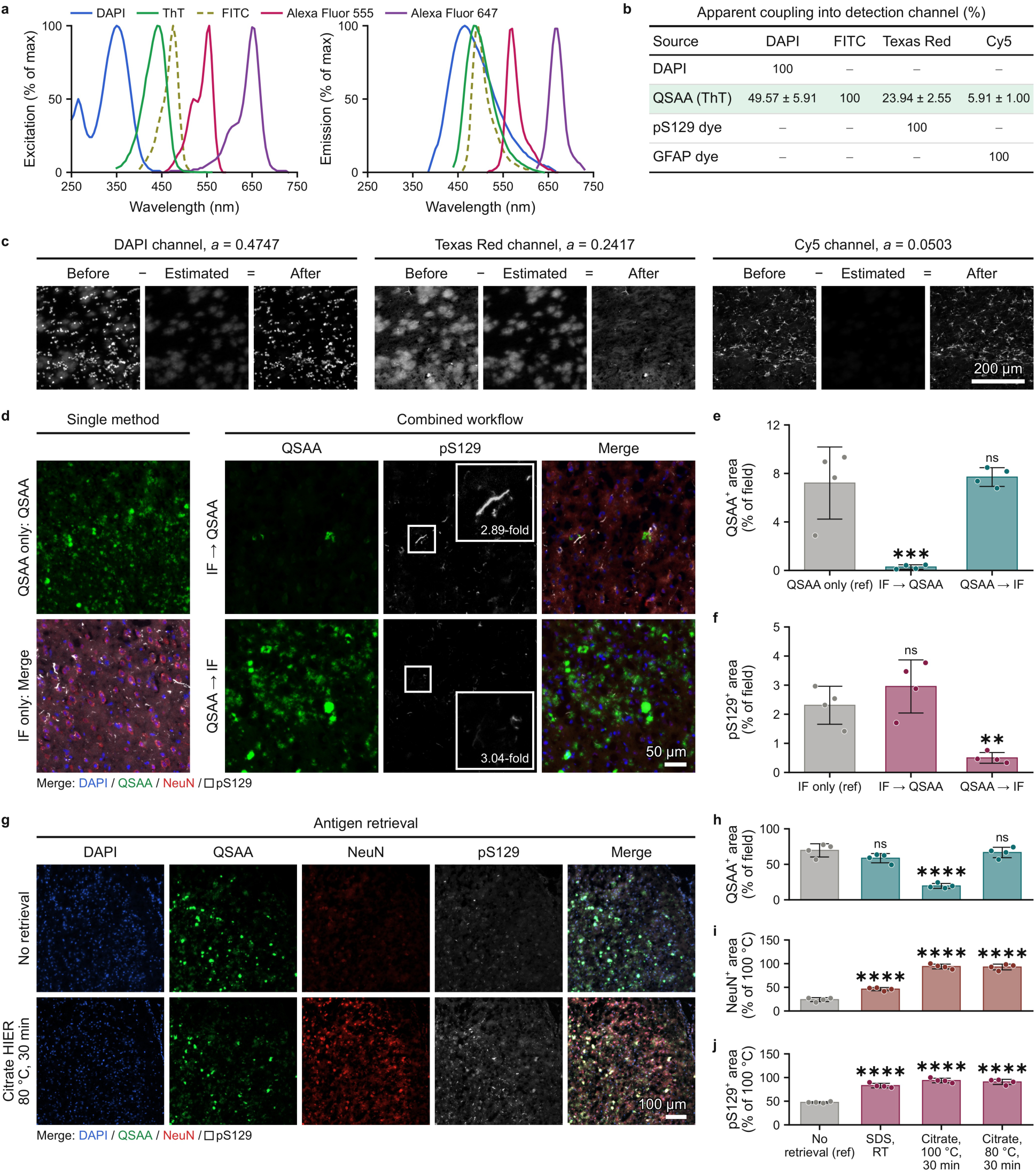
Spectral cross-talk, assay order and antigen retrieval determine how QSAA and immunofluorescence combine on one section. **a**, Excitation (left) and emission (right) spectra of DAPI, ThT, fluorescein isothiocyanate (FITC), Alexa Fluor 555 and 647, each normalized to its maximum. **b**, Apparent coupling of sources (rows) into channels (columns) on a section carrying DAPI, the QSAA product (ThT) and dyes labelling pS129 and GFAP; 100, each source in its own channel; dashes, no detectable coupling. **c**, QSAA component subtracted from DAPI, Texas Red and Cy5 channels of one co-stained field: Before, the channel image; Estimated, the FITC-channel image scaled by header coefficient a; After, their difference. **d**, Sections after QSAA only (top left), IF only (bottom left, merge), IF → QSAA or QSAA → IF. White boxes, inset regions. **e**, **f**, QSAA-positive (QSAA^+^, **e**) and pS129-positive (pS129^+^, **f**) area for workflows in **d**. **g**, Sections without antigen retrieval (top) or citrate heat-induced epitope retrieval (HIER; 80 °C, 30 min; bottom). In merges, DAPI blue, QSAA green, NeuN red and pS129 white (open square in keys). **h**–**j**, QSAA^+^ (**h**), NeuN^+^ (**i**) and pS129^+^ (**j**) area without retrieval, after a sodium dodecyl sulfate (SDS)-based reagent at room temperature (RT) or citrate HIER (80 °C or 100 °C, 30 min); **h**, percentage of field area; **i** and **j**, of signal recovered at 100 °C. Dots, individual values; bars, means; error bars, s.d.; **b**, **e**, **f** and **h**–**j**, n = 4 independent replicates per group. One-way ANOVA with Dunnett’s test against QSAA only (**e**), IF only (**f**) or No retrieval (**h**–**j**), reference group marked (ref); *P < 0.05, **P < 0.01, ***P < 0.001, ****P < 0.0001; ns, not significant. Scale bars, 200 μm (**c**), 50 μm (**d**) and 100 μm (**g**).

The IF → QSAA and QSAA → IF sequences impose a pronounced asymmetric trade-off (Fig. 5d–f). Performing IF prior to QSAA reduced the QSAA-positive area by more than 90% (from 7.2% in QSAA alone to 0.28%; Fig. 5d,e). Even IF without primary antibody caused QSAA product loss exceeding 79–85%, reducing sub-5 μm product density across whole sections from 68 mm^-2^ (No IF) to 10–14 mm^-2^ (Supplementary Fig. 5a–d). Stratification of individual products showed that pre-amplification permeabilization, blocking and overnight washing selectively strip or inactivate small (<5 μm), weakly fluorescent seeds, with losses reaching 92% in the smallest, dimmest stratum while partially sparing large (>10 μm), bright deposits (Supplementary Fig. 5e–h), identifying physicochemical losses during tissue preprocessing as the principal cause of IF → QSAA failure. In solution-phase SAA of seven antibodies, only the conformation-specific MJFR-14-6-4-2 delayed the half-maximal amplification time across all buffer systems (Supplementary Fig. 5i,j). Conversely, performing QSAA first fully preserved the QSAA product area (7.7% vs. 7.2%; Fig. 5e) but reduced pS129 detection roughly fivefold (from 2.31% to 0.50%; Fig. 5f) due to epitope masking during the 70 °C, high-sulfate reaction. Because seed loss during pre-amplification IF is irreversible whereas post-amplification epitope masking is amenable to antigen retrieval, a QSAA-first sequence, stabilized by a 10-min 4% PFA re-fixation step, is mandatory.

To restore antibody accessibility without destroying the newly formed ThT fibril products, we compared five retrieval conditions (Fig. 5g–j). Antibody recovery and product survival stood in direct conflict: citrate heat-induced epitope retrieval (HIER) at 100 °C gave the highest NeuN and pS129 area but the lowest QSAA-positive area, cutting it from 69.7% of field area without retrieval to 19.6% (P < 0.0001; Fig. 5h–j). Across all 12 threshold and object-area combinations, 100 °C heating (citrate or EDTA) thermally disintegrates discrete QSAA fibrils into diffuse background fluorescence, leaving only 0.04–0.23× (citrate 100 °C) and 0.06× (EDTA 100 °C) of the matched no-retrieval control’s discrete product density (Supplementary Fig. 6a–d). Room-temperature SDS reduced discrete product density to 0.35× and recovered NeuN to only 46% of the 100 °C reference (Fig. 5i; Supplementary Fig. 6c,d). By contrast, citrate HIER at 80 °C for 30 min was the only protocol that fully preserved discrete QSAA product density across all 12 threshold combinations (density ratio 1.20×) and left QSAA-positive area statistically unchanged (66.7% versus 69.7%, ns) while restoring NeuN and pS129 immunolabelling (Fig. 5g–j; Supplementary Fig. 6d). This integrated sequence (QSAA → PFA re-fixation → 80 °C citrate HIER → IF → sequential imaging with linear unmixing) constitutes the standardized multiplex IF-QSAA workflow.

### 6. Same-section IF-QSAA reveals associations of seeding activity with neuronal and microglial microenvironments

To apply IF-QSAA to the analysis of cellular microenvironments, and given that neurons and microglia can exhibit distinct αSyn inclusion pathologies [31], we co-stained mPFF-inoculated brain sections for QSAA products alongside neuronal (NeuN), microglial (IBA1) or astrocytic (GFAP) markers and compared the spatial relationships between amplification products and regions labelled by these three markers (Fig. 6a–c). Because amplified QSAA products extend over several micrometres and neurons and glial cells differ substantially in spatial occupancy, raw percentages of touching objects cannot be compared across markers. We therefore coupled IF-QSAA with a spatial permutation null model: within each field, we measured the fraction of QSAA objects contacting marker territory (marker signal occupying at least 5% of a 2-μm ring outside the object boundary) and compared it against 10,000 random rigid translations and rotations within the tissue boundary, preserving object number, size and shape (Fig. 6d).

**Fig. 6.**
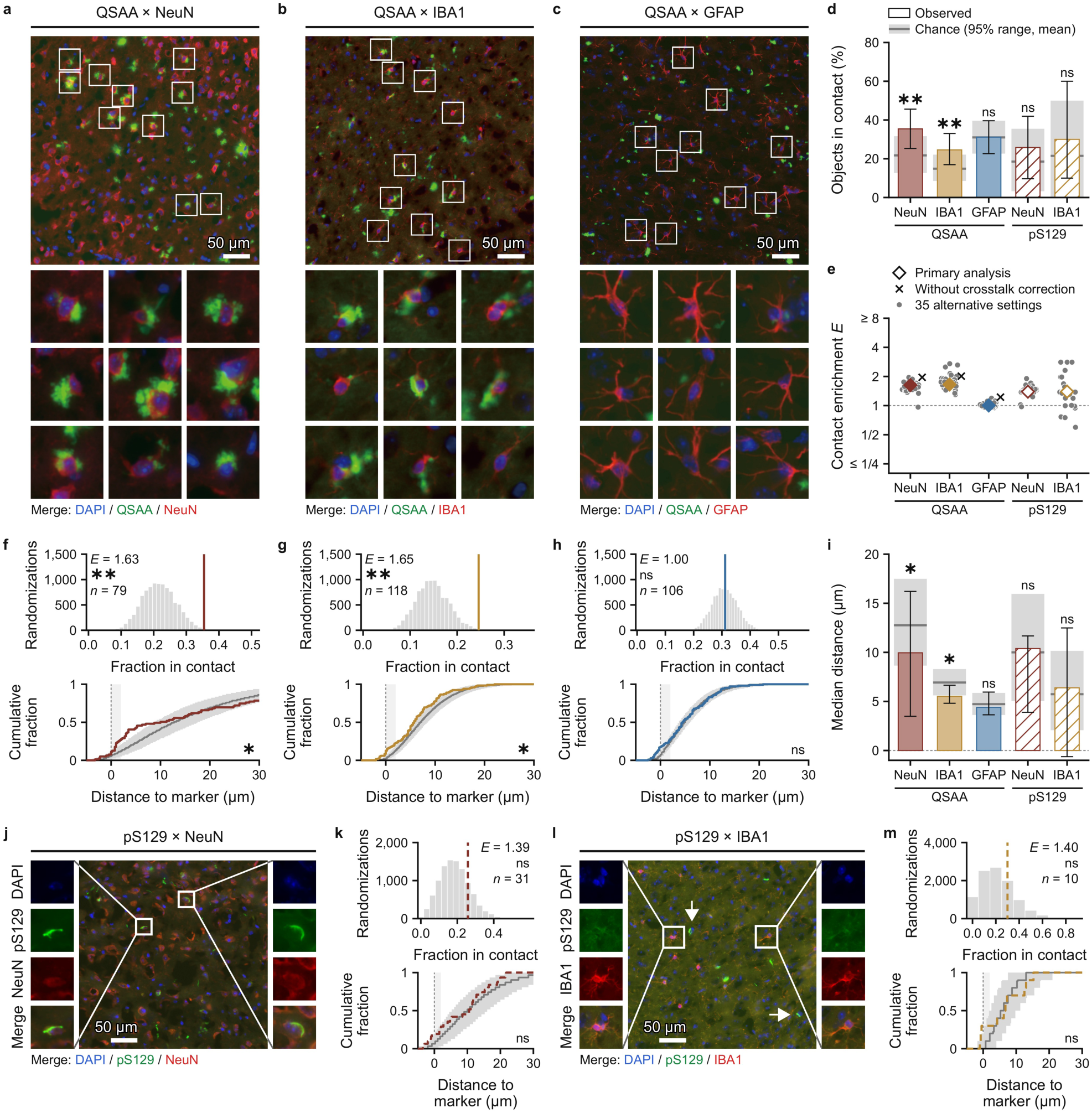
QSAA products contact NeuN- and IBA1-positive cells more often than chance and GFAP-positive cells at the chance rate. **a**–**c**, Brain sections from mPFF-inoculated mice after QSAA and IF for NeuN (neurons; **a**), ionized calcium-binding adapter molecule 1 (IBA1, microglia; **b**) or GFAP (astrocytes; **c**): DAPI, blue; QSAA products, green; marker, red. Boxed regions, enlarged below. **d**, Percentage of QSAA or pS129 objects per field with marker signal within 2 μm of their outline (in contact). Bars, observed; error bars, 2.5–97.5% bootstrap range over objects; grey boxes and lines, 95% range and mean of 10,000 randomizations of object positions (chance). **e**, Contact enrichment E (observed over chance mean): primary analysis (diamonds), no cross-talk correction (crosses; QSAA fields only) and 35 alternative settings (dots); dashed line, chance (E = 1). **f**–**h**, Fields of **a**–**c**. Top, observed fraction in contact (vertical line) against randomizations (histogram); n, objects. Bottom, cumulative distribution of object-centre distance to nearest marker edge (negative, inside): coloured line, observed; grey line and band, randomization median and 95% range; thin vertical line, marker edge; shaded strip, 0–2 μm. **i**, Median distance, as in **d**. **j**–**m**, Conventional IF controls without QSAA: pS129 (green) with NeuN (**j**) or IBA1 (**l**) (red) and DAPI (blue); boxed regions, enlarged as single channels and merge; arrows (**l**), pS129-positive, IBA1-negative profiles. **k**, **m**, These fields as in **f**–**h**, without spectral unmixing (no ThT), only pS129 objects ≥ 100 grey levels. Charts: red, NeuN; ochre, IBA1; blue, GFAP; solid lines, filled bars and symbols, QSAA; dashed lines, hatched bars and open symbols, pS129. Two-sided randomization test (10,000 randomizations) of the fraction in contact (**d**, top) or the whole distance curve (**i**, bottom); P values are not adjusted for multiplicity. *P < 0.05, **P < 0.01, ***P < 0.001, ****P < 0.0001; ns, not significant. Scale bars, 50 μm.

In the co-stained fields (Fig. 6a–c), QSAA products contacted NeuN-positive neuronal regions (35.4% observed vs. 21.7% chance expectation; contact enrichment E = 1.63, P < 0.01, n = 79 objects) and IBA1-positive microglial territories (24.6% observed vs. 14.9% chance; E = 1.65, P < 0.01, n = 118 objects) significantly more often than chance (Fig. 6d,f,g). By contrast, contact with GFAP-positive astrocytic processes matched the randomized null expectation (31.1% observed vs. 31.0% chance; E = 1.00, P = 1.0, n = 106 objects; Fig. 6d,h). Astrocytic processes occupy a wide share of the field area, and nearly one-third of QSAA products touch GFAP by chance alone, yielding an apparent contact rate of 31% and the shortest absolute median object-to-marker distance among the three markers (Fig. 6i), yet an observed contact enrichment close to 1; the randomization null model separates this density-driven contact from genuine spatial association. Studentized envelope tests of the entire cumulative object-to-marker distance curves showed that IBA1 exhibited both significant boundary contact enrichment (E = 1.65) and a significant departure from the randomizations (P = 0.024); NeuN departed over the whole curve as well (P = 0.017), with QSAA products lying clearly denser than random between 0.75 and 7.75 μm from the neuronal boundary and returning to random levels further out (Fig. 6f–i).

Across 35 alternative segmentation and contact parameter settings, contact enrichment remained above 1 for QSAA–IBA1 in all 35 and for QSAA–NeuN in 34, whereas QSAA–GFAP remained near 1 (Fig. 6e). By contrast, in conventional IF controls processed without QSAA, pS129 contact with NeuN and IBA1 did not differ significantly from chance (E = 1.39, P = 0.43, n = 31, Fig. 6j,k; E = 1.40, P = 0.73, n = 10, Fig. 6l,m). The enrichment therefore holds across every threshold tested: amplified seeding activity lies next to neurons and microglia, and astrocytes show no such association.

### 7. Same-section IF-QSAA reveals partial correspondence between seeding activity and pS129 pathology

Same-section IF-QSAA places functional seeding activity and phosphorylated αSyn pathology in a single coordinate system, determining how far the two coincide at the scale of individual microscopic structures. We acquired four-channel IF-QSAA images (DAPI, QSAA, NeuN and pS129) across 75 bilateral fields in 11 mPFF-inoculated mice, sampling five anatomical regions: dorsomedial isocortex (CTX dm), lateral isocortex (CTX lat), caudoputamen (CP), corpus callosum (cc) and hippocampus (HIP) (Fig. 7a; Supplementary Fig. 7a,c). The two readouts displayed distinct morphologies: pS129-positive structures appeared as slender neuritic threads or compact perikaryal inclusions, whereas QSAA products formed larger, rounded deposits (Fig. 7b). This morphological divergence, with abundant single-positive structures, identifies the pS129 signal as endogenous αSyn pathology distinct from the recombinant mouse αSyn monomer recruited into QSAA products.

**Fig. 7.**
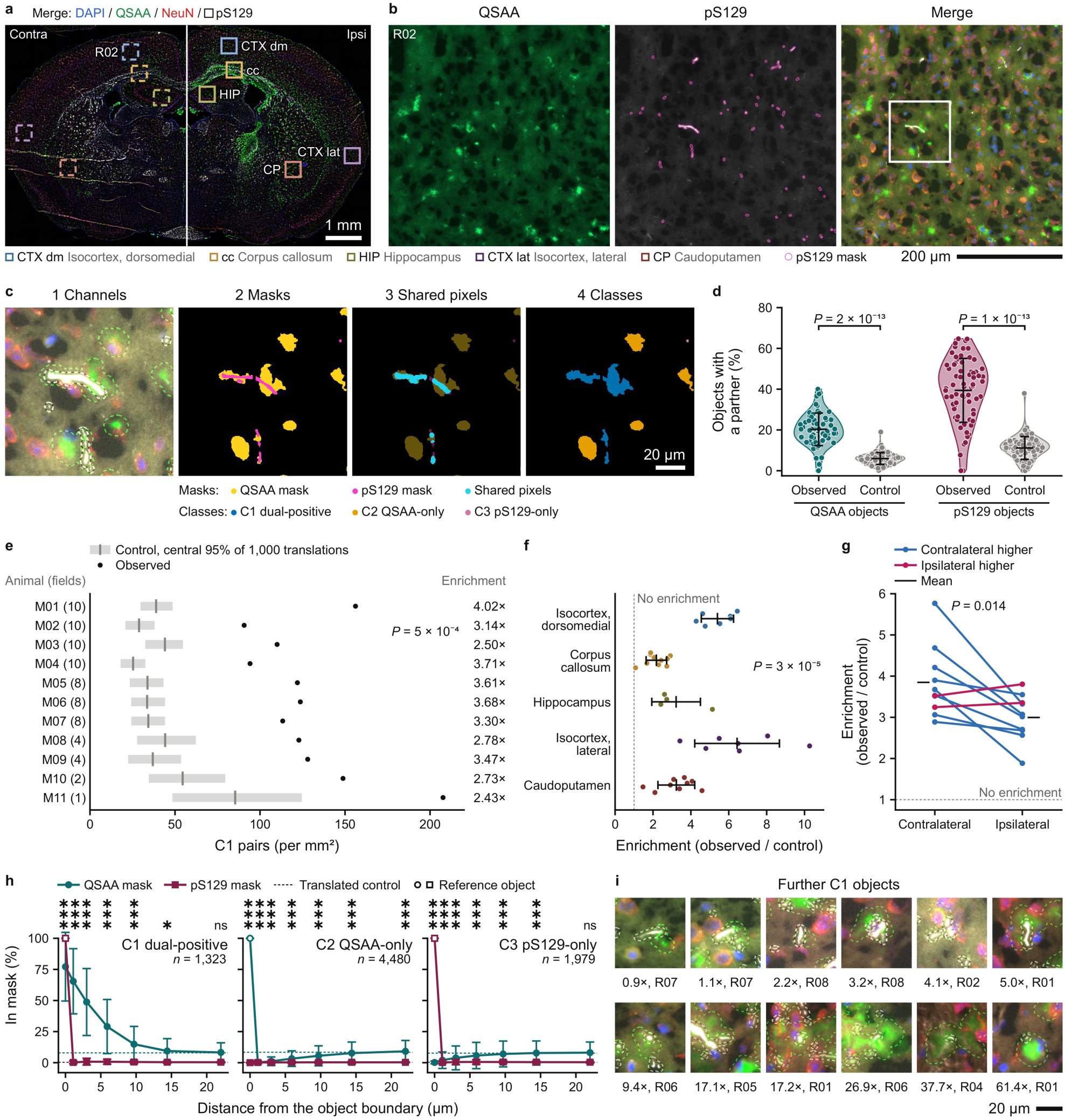
Same-section pairing of QSAA and pS129 objects exceeds chance and varies across brain regions and hemispheres. **a**, Coronal mouse brain section after IF-QSAA: DAPI, blue; QSAA, green; NeuN, red; pS129, white. White line, midline; squares, the ten fields (Supplementary Fig. 7a), solid Ipsi, dashed Contra, colour, region (Supplementary Table 2). **b**, Field R02 enlarged; magenta outlines, pS129 objects counted; white square, **c**. **c**, Classification in the window of **b**: channels with dashed QSAA (green) and pS129 (white) outlines; masks; shared pixels; C1, QSAA–pS129 pairs sharing ≥10% of the smaller object’s area; C2 and C3, unpaired. **d**, Paired objects per field (%), observed and after 1,000 QSAA-mask shifts (control); dots, fields (n = 74, QSAA; 75, pS129); P = 2 × 10^-13^ and 1 × 10^-13^. **e**, C1 pairs per mm^2^ per animal (fields in parentheses); grey tick, control mean; right, enrichment (observed / control); P = 5 × 10^-4^. **f**, Enrichment by region; dots, animals; P = 3 × 10^-5^. **g**, Enrichment on each side; lines, animals imaged on both; P = 0.014. **h**, Percentage of each distance band covered by the other mask, outward from the reference boundary; dashed lines, translated-control whole-field baselines. **i**, Twelve C1 dual-positive pairs centred on the pS129 object, outlined as in **c**; labels, area ratio and field. From 75 fields in 11 animals; Wilcoxon signed-rank test (**d**, **g**, **h**, two-sided; **e**, one-sided); Kruskal– Wallis test (**f**). *P < 0.05, **P < 0.01, ***P < 0.001, ****P < 0.0001; ns, not significant. Scale bars, 1 mm (**a**), 200 μm (**b**) and 20 μm (**c**, **i**).

Using automated pairing, we classified the 5,649 QSAA and 3,302 pS129 objects into three operational categories (Fig. 7c; Supplementary Fig. 7b): C1 dual-positive paired objects, 1,169 QSAA and 1,323 pS129 objects forming 1,332 pairs because one object can overlap several in the other channel; C2 QSAA-only objects (n = 4,480); and C3 pS129-only objects (n = 1,979). Against 1,000 random translations of each field’s QSAA mask, the proportions of QSAA objects (20.7% vs. 6.5%, P = 2 × 10^-13^) and pS129 objects (40.1% vs. 12.2%, P = 1 × 10^-13^) with a partner were above chance (1,332 pairs vs. 402.5 expected, an overall 3.3-fold enrichment; Fig. 7d; Supplementary Fig. 7d). Enrichment was present in all 11 animals (2.43-to 4.02-fold). Animal-level pair densities exceeded translation controls (P = 5 × 10^-4^; Fig. 7e). Enrichment varied by brain region (P = 3 × 10^-5^; Fig. 7f) and was higher in the contralateral than the ipsilateral hemisphere (P = 0.014; Fig. 7g).

Across 38 animal–region summaries, the enrichment ratio correlated negatively with control pair density on logarithmic scales (Pearson r = -0.63; Supplementary Fig. 7f), so high-density regions such as the corpus callosum and ipsilateral hemisphere show lower fold enrichment over a higher random-overlap baseline (Fig. 7f,g).

The object-level census showed that 79.3% of QSAA objects (C2) and 59.9% of pS129 objects (C3) had no counterpart in the other channel (Fig. 7d). Concentric boundary-distance profiling confirmed that C2 objects had zero above-background pS129 coverage within their footprint and across rings to 22 μm (P = 0.77, 0.56 and 0.92 vs. whole-field baseline; Fig. 7h; Supplementary Fig. 7e), proving that they are genuinely unpartnered.

Finally, concentric profiling of paired pS129 objects in C1 (n = 1,323) resolved a core–halo geometry: the QSAA mask covered ∼75–78% of the pS129 object’s interior and extended outward as a halo of elevated coverage over 9.7 μm (P < 0.001 vs. translated control) and 14.4 μm (P < 0.05) before decaying to the whole-field baseline by 22.0 μm (Fig. 7h). This halo scaled monotonically with the area tertile of the QSAA partner (∼20 μm for the largest tertile, 219–1,322 μm^2^; Supplementary Fig. 7g) and recurred in all five anatomical regions at 9.7 μm (Supplementary Fig. 7h). Within individual C1 pairs, the QSAA-to-pS129 area ratio spanned nearly two orders of magnitude, 0.9× to 61.4× (median 17.4×; Fig. 7i), so amplified product area reflects heterogeneous local reaction gain. This geometry traces monomer recruitment and outward extension around pS129-containing seed cores during in situ amplification. Functional seeding activity and Ser129 phosphorylation are statistically associated but non-equivalent pathological states at the level of individual structures.

## Discussion

Building on tissue-section QSAA, we established same-section IF-QSAA to enable direct spatial comparison of αSyn seeding activity with pS129 pathology and cellular markers. This approach advances such comparisons from the regional level to individual pathological structures and reveals a partially overlapping, partially segregated correspondence between seeding signal and pS129 deposition, providing an experimental basis for investigating the molecular state and seeding function of αSyn at the same tissue coordinates.

Such a functional assay is necessary because antigenic detectability and seeding activity are not equivalent features of αSyn pathology. Recombinant PFFs lacking Ser129 phosphorylation already possess the capacity to induce αSyn aggregation, yet cannot be recognized by pS129 antibodies because they lack the corresponding phosphorylated epitope. As a subset of aggregates subsequently acquires this modification, the two signals may transition from segregated to overlapping, consistent with experimental observations that Ser129 phosphorylation can occur after initial aggregation and is further shaped by Lewy body maturation and dynamic post-translational regulation [10,32–35]. Conversely, when IF preprocessing preceded amplification, antibody staining still clearly showed that αSyn pathological deposits remained in the tissue [36], yet the subsequent seeding response dropped markedly, indicating that immunological visibility of pathological material does not guarantee comparable preservation of its detectable seeding competence. This non-equivalence is also evident in spatial distribution: although the two readouts agreed broadly across major grey-matter lesions, several white-matter tracts carried prominent seeding signals without a commensurate degree of pS129 deposition. Together with experimental evidence that αSyn fibrils undergo axonal transport [37], we speculate that these tracts may harbour small particulate seeds that are in transit, not yet extensively phosphorylated, or present at local quantities too low to form immunologically visible deposits. Same-section analysis further showed that dual-positive pairs (C1) occurred at 3.3 times the frequency expected from translated controls—and exceeded all 1,000 translated controls in each of the 11 animals—while substantial populations of QSAA-only (C2) and pS129-only (C3) structures remained. For C2 structures, absence of Ser129 phosphorylation, epitope inaccessibility or insufficient local antigen abundance may each limit immunodetection, whereas seed-templated recruitment of exogenous monomer into abundant new fibrils converts material that is otherwise difficult to observe directly into a visible readout. Thus, QSAA provides not merely higher detection sensitivity, but functional pathological information that cannot be inferred directly from antigenic status.

Applying this functional readout to spatial tissue analysis further requires that the amplified signal be genuinely initiated by tissue-resident seeds and remain traceable to its site of origin. Under our high-reactivity formulation, mechanical agitation accelerated aggregation but also triggered premature unseeded background reactions; removing agitation restored a well-resolved temporal window between seeded and unseeded reactions. The goal of in situ amplification is therefore not simply to maximize reaction speed, but to promote seed-dependent elongation while restraining spontaneous nucleation. At the same time, because QSAA visualizes newly formed fibrils templated by seeds rather than the native pathological material itself, convective migration of seeds or products in the liquid phase would misassign spatial coordinates even in a seed-dependent reaction. Under hydrogel confinement, amplification products expanded progressively while maintaining a median drift-corrected centroid displacement of 1.5–2.0 μm h^-1^—less than a single image pixel (2.78 μm) between consecutive hourly frames—providing a basis for localizing the signal to its originating tissue microdomain. Core–halo profiling of same-section paired objects further showed that QSAA products covered the pS129-positive core and extended into the surrounding periphery, consistent with outward amplification around localized pathological structures and morphologically supporting the reliability of spatial signal assignment. Systematic characterization of seed-dependent amplification kinetics, product localization and their relationship to pre-existing pathological structures thus forms the essential foundation for advancing in situ amplification from simple signal detection to spatial pathological analysis.

Importantly, QSAA measures amplifiability under defined artificial conditions, which is not fully equivalent to recruitment and amplification capacity under physiological conditions in vivo, as the latter is additionally shaped by cellular uptake, proteostatic clearance and local tissue vulnerability. Likewise, detectable templating activity does not equate to the conformational identity of the original seed [38–41]. Using cryo-electron microscopy, Lövestam et al. demonstrated that recombinant αSyn fibrils seeded by multiple system atrophy brain extracts can adopt structures distinct from those of the native filaments [38,39,42], indicating that positive amplification does not guarantee faithful replication of native conformation. Consequently, QSAA reports the presence and magnitude of templating activity under specified reaction conditions, and the original strain cannot be identified from these readouts alone [15,43]. These two considerations define the interpretive boundaries of our measurements with respect to in vivo function and native structure: addressing the former requires correlating IF-QSAA readouts with subsequent pathological burden, changes in cellular function and neuronal injury in combination with interventional studies, whereas addressing the latter requires independent structural comparisons between native tissue filaments and amplified products. Pursuing these directions will also depend on further standardization of the assay across laboratories [19], including defining reaction conditions tailored to sample type and preservation state and standardizing key processing steps and positivity criteria so that procedural variations are not conflated with biological differences.

Future studies may optimize substrates and reaction environments to achieve conformationally faithful in situ amplification, and apply IF-QSAA across disease stages and therapeutic interventions to examine whether changes in pS129 deposition are accompanied by concordant shifts in seeding competence. Extending this framework to other amyloidogenic proteins also has a clear experimental foundation: Saijo et al. and Kraus et al. established RT-QuIC assays for selective tau seeding activity [44,45], while Scialo et al. and Sakamoto et al. developed seed amplification systems for TDP-43 and Aβ [46,47]. These systems provide a starting point for translating protein-specific amplification reactions into localized tissue assays. By establishing appropriate substrates and spatial-confinement conditions, visualizing newly formed aggregates with ThT, and integrating same-section IF to interrogate pathological markers and cellular neighbourhoods, it should be possible to map the seeding activity of multiple disease-associated proteins directly onto specific tissue structures.

## Methods

### Experimental design and study overview

This study was designed to engineer, calibrate and validate multiplexed immunofluorescence-integrated quiescent seed amplification (IF-QSAA) for quantitative spatial mapping of α-synuclein (αSyn) seeding activity in intact brain sections. The methodological framework comprised five sequential stages: (1) biochemical quality control of recombinant αSyn monomers and preformed fibrils (PFFs), followed by solution-phase kinetic engineering of a fragmentation-free, seed-dependent amplification window (Fig. 1 and Supplementary Fig. 1); (2) development of an agarose-hydrogel-confined in situ QSAA protocol on fixed mouse brain sections, validated across coronal, sagittal and horizontal planes and by correlative bright-field and transmission electron microscopy (TEM) (Fig. 2 and Supplementary Fig. 2); (3) whole-brain atlas registration and quantitative regional benchmarking against Ser129-phosphorylated αSyn (pS129) immunofluorescence (IF) and thioflavin S (ThS) staining across grey- and white-matter compartments, early post-inoculation time points and frozen human Parkinson’s disease (PD) brain specimens and archival specimens fixed for over 20 years (Figs. 3 and 4 and Supplementary Figs. 3 and 4); (4) systematic characterization of spectral bleed-through, assay-order-dependent seed depletion and thermal disintegration of fibrils during high-temperature post-amplification antigen retrieval to establish a standardized same-section IF-QSAA workflow (Fig. 5 and Supplementary Figs. 5 and 6); and (5) coupling of same-section IF-QSAA with spatial permutation and translation null models to quantify cellular microenvironment associations (NeuN, IBA1 and GFAP; Fig. 6) and single-object correspondence and core–halo geometry between QSAA products and pS129 pathology (Fig. 7 and Supplementary Fig. 7).

For animal experiments, mice were randomly assigned to experimental groups, and sample sizes were chosen based on prior studies using similar seeding models. Uniform image acquisition settings, segmentation thresholds and quantification procedures were used within each comparison group. The unit of analysis and sample size (n) for each experiment are specified in the corresponding methods and figure legends, distinguishing animals or cases, sections, fields and objects.

### Ethics statement and regulatory oversight

All animal procedures were approved by the Institutional Animal Care and Use Committee (IACUC) of Guangzhou Medical University and were conducted in strict accordance with institutional and national guidelines for the care and use of laboratory animals, in compliance with the ARRIVE guidelines. Post-mortem human brain tissues were obtained from the Department of Neurology, The First Affiliated Hospital of Guangzhou Medical University, with written informed consent from the donors or their next of kin. All procedures involving human tissue were approved by the Ethics Committee of The First Affiliated Hospital of Guangzhou Medical University (approval no. MRE. No. 155) and complied with the Declaration of Helsinki.

### Recombinant αSyn monomer expression and purification

Full-length recombinant mouse wild-type αSyn (mWT, UniProt ID: P58992), human wild-type αSyn (hWT) and human A53T αSyn (hA53T) were expressed in *Escherichia coli* BL21(DE3) cells (TransGen Biotech, CD601) and purified as previously described with modifications [48,49]. Following induction with 1 mM isopropyl β-D-1-thiogalactopyranoside (IPTG) for 4 h at 37 °C, cells were harvested and lysed by sonication in high-salt lysis buffer (10 mM Tris-HCl, pH 7.5, 500 mM NaCl, 1 mM EDTA, 1 mM phenylmethylsulfonyl fluoride [PMSF] and 1× protease inhibitor cocktail [Roche, 04693116001]). The clarified lysate was boiled for 15 min to precipitate heat-labile host proteins, followed by sequential precipitation with 45% saturated ammonium sulfate (Sigma-Aldrich, A4915) and acid precipitation at pH 3.0. Endotoxins were removed using a high-capacity endotoxin removal resin (Sangon Biotech, B641718). The protein was then loaded onto a HiTrap Q HP anion-exchange column (Cytiva, 17115401) equilibrated with 20 mM Tris-HCl (pH 7.5) and eluted with a stepwise NaCl gradient; monomeric αSyn eluted as a single major peak at 30% high-salt buffer B (Supplementary Fig. 1a). Pooled fractions were dialysed against Tris-buffered saline (TBS; 10 mM Tris-HCl, pH 7.5, 50 mM NaCl), passed through a 50-kDa molecular-weight cutoff filter (Amicon Ultra, Millipore, UFC905096) to remove any pre-existing oligomers, and stored at -80 °C. Purity was quantified by Coomassie-stained SDS–PAGE followed by lane densitometry (Supplementary Fig. 1b).

### Preformed fibril (PFF) preparation and characterization

To generate preformed fibrils (mPFFs and hPFFs), purified monomeric αSyn (5 mg mL^-1^ in sterile phosphate-buffered saline [PBS]) was incubated at 37 °C for 7 days with continuous orbital agitation (1,000 r.p.m.) in a ThermoMixer C (Eppendorf, 5382). Fibril suspensions were aliquoted and stored at -80 °C. Immediately prior to inoculation or seeding experiments, fibrils were fragmented using a non-contact water-bath sonicator (SCIENTZ, SCIENTZ08-III) at 90% power for a total of 5 min (300 pulses of 1 s on / 1 s off). Quality control was performed for each PFF batch according to established best-practice guidelines [50]: fibril morphology and length distributions were assessed by negative-stain TEM across five fields per preparation, measuring isolated, complete fibrils (n = 417 mPFFs and n = 21 hPFFs; Supplementary Fig. 1c–e), dynamic light scattering (DLS) confirmed an average hydrodynamic diameter of 50–150 nm, and functional seeding activity was validated in a standard SAA requiring a lag time within ±15% of a reference standard batch when seeded with 1 pg PFFs.

### Solution-phase seed amplification assays and kinetic parameters

#### Cyclic fragmentation (agitated) SAA

Reactions were assembled in 96-well black, clear-bottom plates (Nunc / Thermo Scientific, 165305) in a final volume of 100 μL per well. Unseeded substrate-comparison reactions (Fig. 1c) contained 1 mg mL^-1^ mWT, hWT or hA53T αSyn monomer and 40 μM thioflavin T (ThT; Sigma-Aldrich, T3516). In seeded factor-dissection experiments (Fig. 1d–f), reference control reactions contained 1 mg mL^-1^ hWT αSyn monomer and 40 μM ThT in basal buffer (10 mM Tris-HCl, pH 7.5, 10 mM NaCl) incubated at 37 °C, whereas single-factor and combined variants substituted mWT monomer (1 mg mL^-1^), added 800 mM Na_2_SO_4_ (Sigma-Aldrich, 238597; in 10 mM Tris-HCl, pH 6.0, 25 mM NaCl), or elevated the incubation temperature to 70 °C as indicated. Seeded wells received 1 pg PFFs; unseeded control wells received an equal volume of buffer. Plates were sealed with optical adhesive film (Applied Biosystems, 4311971) and incubated in a fluorescence plate reader (SpectraMax iD5, Molecular Devices) with recurrent cycles of 1 min double-orbital shaking (700 r.p.m.) followed by 14 min rest. Bottom-read ThT fluorescence (excitation 450 nm, emission 490 nm) was recorded every 15 min.

#### Quiescent SAA

Reactions were prepared in 96-well polypropylene PCR plates (Thermo Scientific, AB0700) in a final volume of 20 μL containing 1 mg mL^-1^ mWT αSyn monomer (Fig. 1g) and 40 μM ThT in QSAA reaction buffer (800 mM Na_2_SO_4_, 25 mM NaCl, 10 mM Tris-HCl, pH 6.0), seeded with 1 pg mPFFs or an equal volume of buffer as an unseeded control. Each well was overlaid with 20 μL mineral oil (Biosharp, BS927) and sealed to prevent evaporation at high temperature. Plates were incubated at 70 °C without any shaking or sonication in a real-time fluorescence thermal cycler (Bio-Rad CFX96) or fluorescence reader, with ThT fluorescence recorded hourly or every 30 min via the FAM/FITC channel (excitation 452–486 nm, emission 505–545 nm).

#### Kinetic metrics

The lag phase (h) was defined as the time required for ThT fluorescence to cross the positivity threshold (baseline mean plus 5 standard deviations); wells that remained at baseline throughout the observation window were assigned the run ceiling (100 h) for conservative statistical comparison. The protein aggregation rate (PAR, h^-1^) was calculated as the reciprocal of the lag phase (1 / lag phase). The dimensionless PAR ratio was defined as the mean seeded PAR divided by the mean unseeded PAR under the same reaction condition.

### Animals and stereotaxic inoculation

Male C57BL/6J mice (10–12 weeks old; RRID: IMSR_JAX:000664) were obtained from the Experimental Animal Center of Guangzhou Medical University and housed under specific-pathogen-free (SPF) conditions on a 12-h light/dark cycle with *ad libitum* access to food and water. For stereotaxic surgery, mice were anaesthetized with isoflurane (2.5% induction, 1.5% maintenance) and positioned in a stereotaxic frame (Kopf Instruments, Model 940). Mice received a unilateral injection into the right dorsal striatum at coordinates relative to Bregma (Franklin and Paxinos atlas, 3rd edition): anteroposterior (AP) +0.8 mm, mediolateral (ML) +2.0 mm, and dorsoventral (DV) -3.5 mm. A total of 5 μg sonicated mPFFs in 1 μL PBS or 1 μL sterile PBS (vehicle control) was infused at 0.2 μL min^-1^ using a 10-μL Hamilton syringe fitted with a 33-gauge needle (Hamilton, 65460-05) driven by a microinfusion pump. The needle was left in place for 10 min post-infusion to prevent reflux before slow withdrawal.

At designated endpoints (7, 10, 14, 30 or 180 days post-inoculation [dpi]), mice were deeply anaesthetized and transcardially perfused with ice-cold PBS followed by 4% (w/v) paraformaldehyde (PFA) in 0.1 M phosphate buffer (pH 7.4). Brains were post-fixed in 4% PFA at 4 °C for 24 h, cryoprotected in 30% (w/v) sucrose in PBS until equilibrated, embedded and cryosectioned at 14 μm thickness onto high-adhesion glass slides (CITOTEST, 188105W).

### Post-mortem human brain specimens and pretreatment of long-term fixed tissue

Post-mortem human brain tissues (collected between 2005 and 2022) were obtained from The First Affiliated Hospital of Guangzhou Medical University from neuropathologically confirmed cases of Parkinson’s disease (PD, n = 7), progressive supranuclear palsy (PSP, n = 6) and age-matched neurologically normal healthy controls (HC, n = 6) with post-mortem intervals < 12 h.

Neuropathological diagnoses were established based on consensus criteria, including Braak staging for Lewy body pathology. For standard cases, one hemisphere was fixed in 10% neutral buffered formalin (NBF) for 2–4 weeks while the contralateral hemisphere was coronally sliced and snap-frozen at -80 °C. In addition to standard samples, an archival PD brain specimen that had been stored in 10% NBF at room temperature for more than 20 years (FF-PD, four technical replicates) was examined to evaluate the feasibility of in situ amplification after pretreatment of long-term fixed archival tissue. Prior to cryosectioning at 14 μm, FF-PD tissue blocks were incubated in PBS at 95 °C for 1 h to partially reverse formalin cross-links, followed by equilibration in 30% sucrose. Regional human comparisons sampled five anatomical sites: corpus callosum (CC), two striatal sites (Striatum (L) and Striatum (R)) and two cortical sites (Cortex (L) and Cortex (R)) (Supplementary Fig. 4e,f).

### In situ QSAA on brain sections

Cryosections (14 μm) mounted on adhesion slides were air-dried for 30 min, post-fixed in 4% PFA for 10 min at room temperature, washed in PBS and air-dried. To confine the amplification reaction and suppress micro-convection over the tissue surface, a pre-warmed (∼ 50 °C) 1% (w/v) aqueous agarose solution (BioFroxx, 1110GR100; in ddH_2_O) was cast over the section to form a uniform hydrogel layer approximately 1 mm thick. After gelation at room temperature for 5 min, excess agarose outside the tissue perimeter was trimmed flush with a scalpel.

The QSAA reaction mixture was freshly prepared by combining equal volumes (150 μL : 150 μL) of a 2× buffer stock (∼ 3.41 g Na_2_SO_4_, 1.2 mL of 1 mM ThT and 150 μL of 1 M Tris-HCl pH 5.5 in 15 mL water) and a 2× substrate stock (2 mg mL^-1^ mWT αSyn monomer in 10 mM Tris-HCl pH 7.5, 50 mM NaCl), yielding final working concentrations of 1 mg mL^-1^ αSyn monomer, ∼ 800 mM sodium sulfate, 10 mM Tris-HCl (pH 6.0), 25 mM NaCl and 40 μM ThT. A 300-μL aliquot of reaction mixture was applied over the agarose-coated section, and slides were incubated quiescently in a sealed humidified chamber on a precision temperature-controlled stage (Ruicheng Instrument Co., TS400) at 70 °C for 8 h. After incubation, the agarose overlay was gently floated off in pre-warmed PBST, and sections were washed in PBST for 1 min before mounting or downstream processing. For real-time in situ tracking (Supplementary Fig. 4g,h), human PD sections undergoing QSAA at 70 °C were imaged at 0, 1, 2, 3, 4 and 5 h across a fixed region of interest.

### Transmission electron microscopy (TEM)

For solution-phase quiescent amplification products (Supplementary Fig. 1c,f–i), 10 μL of reaction mixture or sonicated PFF suspension was adsorbed onto glow-discharged 400-mesh carbon-coated copper grids for 2 min, rinsed with ultrapure water and negatively stained with 2% (w/v) uranyl acetate. For in situ ultrastructural analysis of reacted brain tissue (Fig. 2d), QSAA-reacted sections were carefully detached from slides, fixed in 2.5% glutaraldehyde and 2% PFA in 0.1 M sodium cacodylate buffer (pH 7.4), post-fixed in 1% osmium tetroxide, dehydrated through a graded ethanol series and embedded in EMBed 812 resin. Ultrathin sections (70 nm) were cut on an ultramicrotome, collected on copper grids, stained with uranyl acetate and lead citrate, and imaged on a FEI Talos L120C transmission electron microscope operating at 120 kV.

### Orthogonal validation by brain homogenate RT-QuIC

Human brain tissues (PD, FF-PD, PSP and HC) were homogenized at 10% (w/v) in ice-cold PBS using a bead beater, clarified by brief centrifugation and serially diluted (10^-2^ to 10^-5^ in PBS). Reactions were performed in 96-well plates containing 100 μL of RT-QuIC reaction mix (0.1 mg mL^-1^ recombinant human αSyn [1–140], 10 μM ThT, 10 mM Tris-HCl pH 7.5 and 50 mM NaCl) seeded with 2 μL of diluted brain homogenate (n = 3 replicate reactions per group). Plates were incubated at 37 °C in a fluorescence plate reader (BMG FLUOstar Omega) with cycles of 1 min double-orbital shaking (700 r.p.m.) and 1 min rest for 16 h, recording ThT fluorescence (excitation 450 nm, emission 490 nm) every 15 min (Supplementary Fig. 4c,d). A sample was scored positive when at least two of three replicates crossed the threshold of baseline mean plus 5 standard deviations.

### Multiplexed same-section IF-QSAA workflow and optimization

#### Assay-order evaluation

To evaluate procedural interference between QSAA and IF (Fig. 5d–f and Supplementary Fig. 5), brain sections were processed under four regimes: (i) standalone QSAA; (ii) standalone IF; (iii) IF → QSAA, in which sections underwent standard IF processing (permeabilization, blocking, overnight incubation at 4 °C without primary antibody or with anti-NeuN + anti-Olig2, EP1536Y, MJFR-14-6-4-2 or 4D6, and washing) prior to QSAA; and (iv) QSAA → IF, in which QSAA was performed prior to IF. The tissue-pretreatment comparisons were repeated in four independent experimental batches, with spatial sampling by image tile within each section. To test direct antibody effects on solution kinetics (Supplementary Fig. 5i,j), seven antibodies (1:100 dilution, corresponding to a tenfold higher concentration than that used for tissue IF with the same antibody) were added to plate-based SAA reactions in ammonium sulfate (AS) or sodium sulfate (SS) buffers with human or mouse substrates, and the shift in half-maximal fluorescence time (Δ T1/2, in cycles of ∼ 31 min) was quantified relative to no-antibody controls.

#### Post-amplification fixation and antigen-retrieval calibration

In the standardized QSAA-first workflow, freshly cut 14-μm sections were fixed in 4% PFA for 10 min, washed in PBS and subjected to QSAA for 8 h. Immediately after removing the agarose overlay, sections were re-fixed in 4% PFA for 10 min at room temperature to cross-link and stabilize the newly formed amplification products against subsequent thermal and detergent treatments. We then compared five post-QSAA antigen-retrieval conditions prior to immunostaining (Fig. 5g–j and Supplementary Fig. 6a–d): (1) no retrieval; (2) an SDS-based antigen retrieval solution at room temperature (P0090, Beyotime); (3) 1 mM EDTA buffer (pH 8.0) at 100 °C for 30 min; (4) 10 mM sodium citrate buffer (pH 6.0) at 100 °C for 30 min; and (5) 10 mM sodium citrate buffer (pH 6.0) at 80 °C for 30 min (the adopted optimal condition).

#### Immunostaining and Thioflavin S staining

Following 80 °C citrate retrieval and cooling to room temperature, sections were permeabilized with 0.5% Triton X-100 in PBS for 15 min and blocked with 5% normal goat serum (or 5% bovine serum albumin) in PBS for 1 h at room temperature. Sections were incubated overnight at 4 °C with primary antibodies (all at 1:1,000 dilution; Supplementary Table 1) against pS129-αSyn (rabbit monoclonal clone EP1536Y, Abcam, ab51253; or mouse monoclonal clone 81A, Abcam, ab184674), NeuN (mouse monoclonal clone A60, Millipore, MAB377), IBA1 (rabbit polyclonal, FUJIFILM Wako, 019-19741) or GFAP (mouse monoclonal, Cell Signaling Technology, 3670). After three PBS washes, sections were incubated for 1 h at room temperature with species-appropriate secondary antibodies (1:1,000, Invitrogen; Supplementary Table 1) conjugated to Alexa Fluor 555 (goat anti-rabbit IgG, A-21428; goat anti-mouse IgG, A-21422; read in the Texas Red channel) and/or Alexa Fluor 647 (goat anti-rabbit IgG, A-21245; goat anti-mouse IgG, A-21235; read in the Cy5 channel), counterstained with 4′,6-diamidino-2-phenylindole (DAPI) and mounted with antifade medium. In standalone IF controls, the corresponding secondary antibodies were conjugated to Alexa Fluor 488 (goat anti-rabbit IgG, A-11008; goat anti-mouse IgG, A-11001). For ThS comparison (Fig. 4a,b), sections were incubated in 0.05% ThS (Sigma-Aldrich, T1892) in 50% ethanol for 8 min, differentiated in 80% ethanol, treated with TrueBlack lipofuscin autofluorescence quencher (Biotium, 23007) and mounted.

All antibodies, chemicals and buffers, consumables and columns, instruments and software used in this study are listed in Supplementary Table 1.

### Multichannel microscopy and linear spectral unmixing

Whole-section scans and fluorescence microfields were acquired on a Nikon Eclipse Ti2-E widefield fluorescence microscope controlled by NIS-Elements Advanced Research software (v5.30, Nikon; 4×, 10× and 20× S Plan Fluor ELWD objectives, Digital Sight 50M camera). All images within each comparison group were acquired with identical exposure, detector gain and offset settings.

To reduce cross-excitation during multiplexed imaging, channels were acquired sequentially in descending order of wavelength: Cy5/Alexa Fluor 647 (excitation 633 nm, emission 650–720 nm) → Texas Red/Alexa Fluor 555 (excitation 561 nm, emission 570–630 nm) → FITC/ThT (QSAA) (excitation 488 nm, emission 495–550 nm) → DAPI (excitation 405 nm, emission 415–470 nm). To account for ThT signal extending into adjacent detection channels (Fig. 5a), apparent channel-coupling coefficients were calibrated on single-source and co-stained sections (n = 4; Fig. 5b). Prior to segmentation and quantitative spatial analysis, pixel-wise linear spectral unmixing was applied to each non-FITC channel c ∈ {DAPI, Texas Red, Cy5} according to:

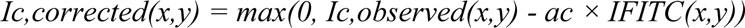

where the empirically calibrated coupling coefficients were aDAPI = 0.4747, aTexas Red = 0.2417 and aCy5 = 0.0503 (Fig. 5c). This subtraction model was applied to ThT-containing same-section IF-QSAA images in Figs. 5 and 7 and Supplementary Figs. 6 and 7; cross-talk handling for Fig. 6 is described under ’Cellular contact and spatial permutation null model analysis’.

### Whole-brain atlas registration and regional quantification

Whole-section QSAA and matched pS129 IF coronal scans at six rostro-caudal levels (Bregma+1.46, +1.11, -0.92, -1.51, -3.07 and -4.54 mm) were registered to the Allen Mouse Brain Reference Atlas (and annotated alongside Franklin and Paxinos nomenclature). Registration accuracy was evaluated using the Dice similarity coefficient between the segmented tissue outline and the corresponding atlas plate, yielding Dice values of 0.90 (+1.46 mm), 0.88 (+1.11 mm), 0.84 (-0.92 mm), 0.83 (-1.51 mm), 0.88 (-3.07 mm) and 0.53 (-4.54 mm) (Fig. 3a and Supplementary Fig. 3a– e). Because registration at Bregma -4.54 mm was unreliable (Dice 0.53), regional share maps and division-level summaries (Fig. 3c,f,g) were restricted to the five well-registered levels (+1.46 to - 3.07 mm).

Segmented objects (4,459 QSAA objects and 3,400 pS129 objects across the five levels, after excluding fold and cut-edge artefacts and the confluent injection-site mass >1 mm^2^ so as not to skew remote regional shares) were apportioned to atlas regions according to their spatial footprint. Regional distributions in Fig. 3c,f were normalized as each region’s percentage share of that marker’s own total object area across the five levels. For white-matter tract enrichment (Fig. 3g), a tract’s share of footprint-weighted QSAA objects was divided by its share of pS129 objects (normalized to the hemispheric baseline ratio). Uncertainty and stability were assessed by 95% bootstrap confidence intervals and leave-one-level-out sensitivity ranges; tracts were classified as robust (filled symbols) only if they contained at least five whole objects of each marker and both the 95% bootstrap CI and the leave-one-level-out range excluded 1.0.

### Object segmentation, multi-threshold retrieval grid and time-lapse tracking

Image segmentation and object feature extraction were performed using custom Python pipelines (incorporating FluoroAnts v6.3, OpenCV and scikit-image) alongside Fiji/ImageJ (v2.3.0, NIH). For the post-QSAA antigen-retrieval calibration in Supplementary Fig. 6d, discrete QSAA products were defined as 8-connected components brighter than the local background median (computed within a 40-μm sliding window) by a specified intensity threshold and exceeding a minimum object area. To ensure that conclusions regarding thermal product destruction were independent of arbitrary thresholding, product density (objects mm^-2^, normalized to the matched no-retrieval control) was evaluated across a 12-parameter grid combining four local intensity thresholds (15, 25, 40 and 60 grey levels above local background) and three minimum object areas (3.5, 7 and 14 μm^2^).

For the time-lapse growth analysis in human PD tissue (Supplementary Fig. 4g,h), frames acquired from 0 to 5 h were aligned by phase correlation of tissue texture to correct for stage drift, objects were segmented at a fixed threshold within a 1.78-mm (640 × 640 px) ROI, and one-to-one matched objects between consecutive hourly frames (n = 6, 29, 70, 98 and 63 pairs across the five intervals) were used to compute median hourly centroid displacement and its interquartile range.

### Cellular contact and spatial permutation null model analysis

We conducted the cellular microenvironment experiments (Fig. 6) in three independent batches using different animals, and we selected representative fields for object-level spatial analysis. Background was subtracted using a 19-μm rolling ball, and candidate QSAA or pS129 objects (minimum area 0.5 μm^2^) and cellular marker territories (NeuN, IBA1 or GFAP) were segmented into binary masks. Objects were thresholded at the larger of the background mean plus four noise standard deviations and a grey-level floor; the floor was 51 in the ThT-containing QSAA fields (Fig. 6a–c) and 100 (8-bit scale) in the ThT-free conventional IF fields (Fig. 6j,l), so that only bright pS129 signal was counted there. In ThT-containing fields, red–green cross-talk in both directions and DAPI-to-readout cross-talk were estimated per field from the image and removed by linear unmixing before segmentation; ThT-free fields were not unmixed. In all fields, readout bleed-through into the DAPI channel was subtracted before nuclei were masked, marker components lying ≥ 80% within the readout object footprint dilated by 1 μm were removed, and objects whose readout contrast against a surrounding 1–3-μm ring was lower than the DAPI or marker contrast were excluded. Operational contact was scored when cellular marker territory occupied ≥ 5% of the 2-μm dilation ring immediately outside an object’s boundary. Signed object-to-marker distance was measured from each object’s centroid to the nearest edge of the cellular marker mask (negative values indicating centroids inside the marker mask).

To eliminate chance-contact bias arising from differences in marker density and amplified product size, a spatial null distribution was constructed within each field. Object number, area and shape were held constant while all objects underwent 10,000 random rigid transformations within the valid tissue boundary. Transformations combined rotations in multiples of 90° with toroidal translations >20 μm; invalid transformations entering tissue voids were automatically redrawn. Contact enrichment (E) was defined as the observed contact fraction divided by the mean contact fraction across the 10,000 randomizations:

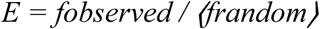

where fobserved is the observed fraction of objects in contact and ⟨frandom⟩ is the mean contact fraction across the 10,000 randomizations. Two-sided empirical P values were computed from the randomization distribution without adjustment for multiplicity. The distance facets were tested over the whole curve rather than at the median: the statistic is the largest studentized departure of the observed cumulative distribution from the randomization mean, sup|ECDF − ⟨ECDFrandom⟩| / s.d.random, evaluated over −5 to 30 μm. Analytical robustness was tested across 35 alternative parameter settings varying object thresholds, marker smoothing/closing and contact ring widths (Fig. 6e).

### Same-section object pairing and concentric boundary-distance profiling

In the same-section IF-QSAA dataset (75 fields of 418 × 418 μm across five bilateral brain regions from 11 mice; Fig. 7 and Supplementary Fig. 7), binary masks of QSAA products and pS129 inclusions were generated independently. A QSAA object and a pS129 object were classified as a C1 dual-positive pair when their masks shared ≥ 10% of the area of the smaller object; all remaining objects were classified as C2 (QSAA-only) or C3 (pS129-only) (Fig. 7c).

To test whether observed pairing exceeded chance overlap determined by local object densities, the QSAA mask of each field was spatially translated relative to the pS129 mask by 40– 200 pixels (15–76 μm) 1,000 times within the frame, recounting paired objects on each shift. Enrichment was calculated as the observed pair count (or density) divided by the mean of the 1,000 translated controls at the field, regional, hemispheric and animal levels.

To characterize the local spatial geometry around individual objects (Fig. 7h and Supplementary Fig. 7e,g,h), concentric distance bands (annuli extending to 22.0 μm, evaluated at midpoint distances including 9.7, 14.4 and 22.0 μm) were constructed outward from the boundary of each reference object (pS129 mask for C1 and C3; QSAA mask for C2), and the percentage of each annulus covered by the second mask was compared against the whole-field translated baseline using paired tests.

### Statistical analysis

Statistical analyses were performed in Python (NumPy, SciPy and pandas) and GraphPad Prism (v10.2.2). Prior to applying parametric tests, data distributions were assessed for normality using the Shapiro–Wilk test and for homogeneity of variances using Levene’s test. Multi-group comparisons with a single control group used one-way analysis of variance (ANOVA) followed by Dunnett’s post-hoc test (Fig. 1c,e, Fig. 5e,f,h–j and Supplementary Fig. 4b); all-pairwise multi-group comparisons used one-way ANOVA followed by Tukey’s test (Fig. 4b). Two-group comparisons used two-tailed unpaired Student’s *t*-tests, applying Welch’s correction for unequal variances and Holm’s step-down adjustment for multiple comparisons across time points or anatomical levels where indicated (Fig. 1g, Fig. 4d,f and Supplementary Figs. 4f and 5j). Cellular contact within fields was evaluated by two-sided randomization tests (10,000 randomizations); the distance distributions were tested over the whole curve by a studentized envelope test on the same randomizations. Neither was adjusted for multiplicity (Fig. 6). Paired comparisons of observed versus translation-control pairing across fields and animals used exact Wilcoxon signed-rank tests (Fig. 7d,e,g,h and Supplementary Fig. 7e), and multi-region enrichment comparisons used the non-parametric Kruskal–Wallis / exact Friedman test (Fig. 7f). Data are presented as mean ± s.d. or median with interquartile range (or 95% bootstrap CIs) as specified in each legend.

## Data Availability

Full-resolution micrographs, whole-section scans, segmentation masks and plate-reader kinetic exports are available from the corresponding author on request. Anatomical reference plates were adapted from the publicly available Allen Mouse Brain Common Coordinate Framework (CCFv3) and the Franklin & Paxinos Mouse Brain Atlas.

## Code Availability

The custom Python scripts used for spectral unmixing, object segmentation (including FluoroAnts v6.3), whole-brain atlas registration, 10,000-draw spatial permutation contact analysis, 1,000-shift mask translation controls, concentric annulus profiling and figure compilation are available from the corresponding author on request.

## Acknowledgements and Funding

We thank the brain tissue donors and their families for their generous contribution to medical research, and the staff of the Department of Neurology and Department of Pathology at The First Affiliated Hospital of Guangzhou Medical University for assistance with neuropathological characterization and electron microscopy. This work was supported by the National Natural Science Foundation of China (grant nos. U25A2059 and 82071416), the Guangdong Basic and Applied Basic Research Foundation (grant no. 2022B1515230004), the Xinjiang Uygur Autonomous Region Key Research and Development Program (grant no. 2023B03003), the Guangzhou Municipal Key Research and Development Program (grant no. 2023B03J0631), the 2024 Basic Research Program of City–University (Hospital) Joint Funding ‘Dengfeng Hospital’ of the Guangzhou Municipal Science and Technology Bureau (grant no. 2023A03J01331) and the Clinical High-Tech Project of the Guangzhou Municipal Health Commission (grant no. 2026C-GX009). The funders had no role in study design, data collection and analysis, decision to publish or preparation of the manuscript.

## Author Contributions

H.M., Y.K. and P.-Y.X. conceived and designed the study. H.M. and Y.K. developed the QSAA and multiplex IF-QSAA methodologies, performed the biochemical, animal and human tissue experiments, and acquired the microscopy data. W.D., H.L., T.G. and Z.H. assisted with stereotaxic surgeries, tissue processing and reagent preparation. J.L., C.M. and J.H. established the animal models and carried out the associated analyses and statistics. H.M. and Y.K. developed the quantitative image-analysis pipelines and performed the statistical analyses. W.-Q.Z., Y.H., Z.W. and P.-Y.X. provided resources, neuropathological expertise and project supervision. H.M., Y.K., Y.H., Z.W. and P.-Y.X. wrote and revised the manuscript. All authors reviewed and approved the final manuscript.

