## Supplementary Material for "Spatial mapping of α-synuclein seeding activity and pathology using immunofluorescence-integrated quiescent seed amplification"

Supplementary Figures 1–7 and Supplementary Tables 1 and 2.

### Supplementary Figures

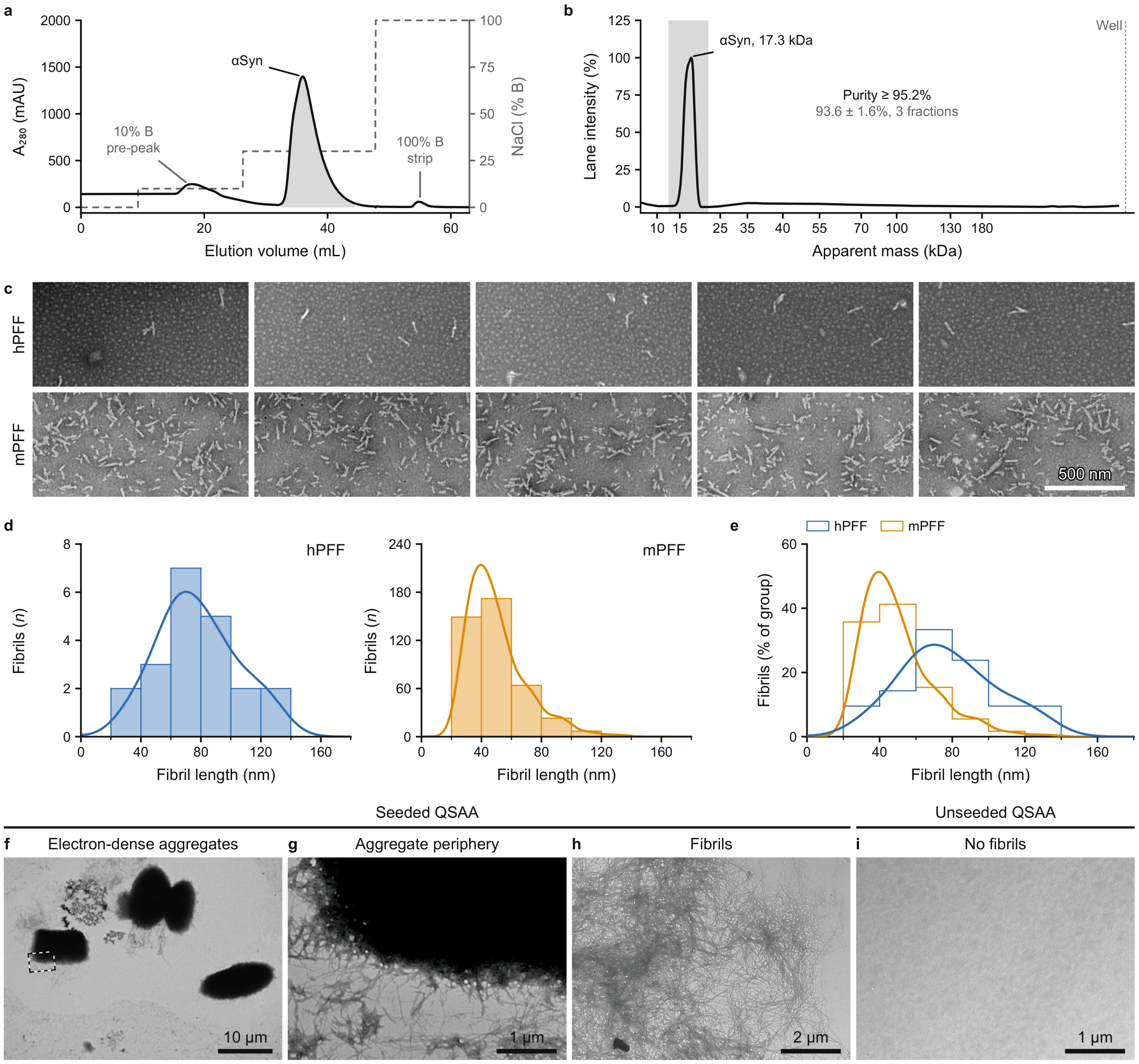


**Supplementary Fig. 1 | Recombinant α-synuclein is purified by ion-exchange chromatography, both preformed fibril preparations consist of short fibrils, and the quiescent amplification products contain fibrils.** **a**, Ion-exchange chromatogram of recombinant α-synuclein (αSyn) eluted with a stepwise NaCl gradient. Black line, absorbance at 280 nm in milli-absorbance units ($A_{280}$, mAU; left axis); grey dashed line, high-salt buffer B percentage (right axis); grey shading, the αSyn peak collected at 30% B; grey labels, the minor peaks at 10% B and the 100% B column strip. **b**, Densitometric profile of one SDS–PAGE lane loaded with a 30% B fraction. Black line, intensity as a percentage of the lane maximum; x axis, lane position, labelled with the apparent masses of the marker bands; grey band, the αSyn band; dotted line, loading well; black text, purity of this lane; grey text, purity of three fractions. **c**, Electron micrographs of human (hPFF, top row) and mouse (mPFF, bottom row) αSyn preformed fibrils, five fields each. **d**, Lengths of individual fibrils measured in **c**, for hPFF (blue, left) and mPFF (orange, right). Bars, fibril counts in 20 nm bins; curves, kernel density estimates. **e**, The same lengths as a percentage of fibrils in each group; step outlines, histograms; curves and colours as in **d**. **f**–**i**, Transmission electron micrographs of seeded (**f**–**h**) and unseeded (**i**) QSAA reaction products; the product takes two forms. **f**, Electron-dense aggregates at low magnification. **g**, The boxed region in **f** enlarged; fibrils at the aggregate periphery. **h**, Fibrils, the second form. **i**, No fibrils. In **b**, mean ± s.d.; n = 3 fractions; in **d** and **e**, n is the number of fibrils (hPFF, 21; mPFF, 417). Scale bars, 500 nm (**c**), 10 μm (**f**), 1 μm (**g**, **i**) and 2 μm (**h**).

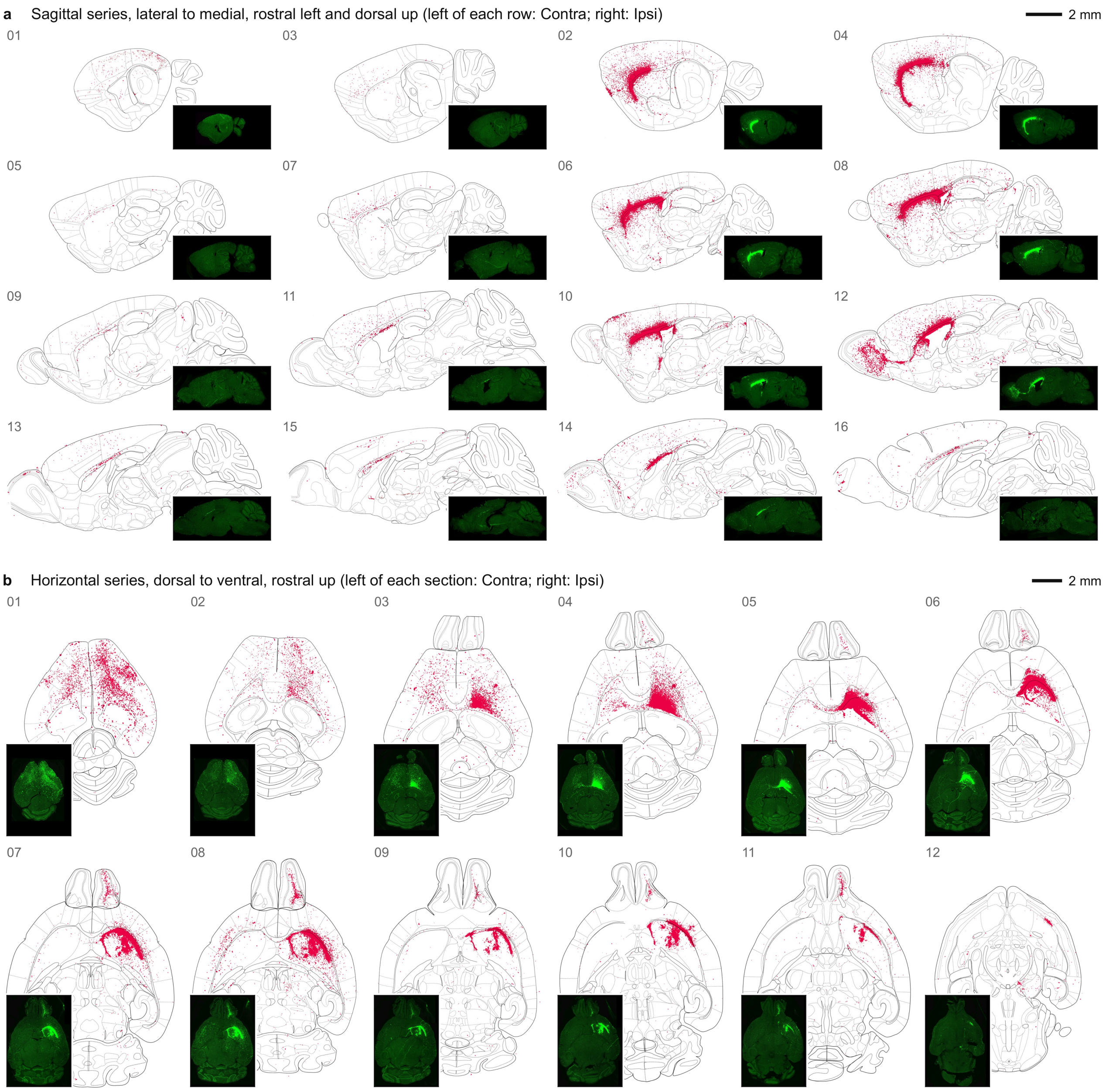


**Supplementary Fig. 2 | Object maps of sagittal and horizontal section series show where the QSAA signal lies across the whole mPFF-inoculated mouse brain.** **a**, **b**, QSAA objects of whole sections of mouse brain, sixteen of the sagittal (**a**) and twelve of the horizontal (**b**) series, 180 days after inoculation of mPFF into one striatum. Each section is shown as its atlas map (grey outlines): every QSAA object is a crimson dot, all sections of a panel in the same frame and at the same scale. The inset on each map is the ThT fluorescence of that section (green), the image the map was made from, at about a third of the map's width (bottom right in **a**, bottom left in **b**, whichever corner covers the fewest objects). All insets use a uniform crop frame around the tissue and a monotonic gamma adjustment (0.65), without clipping values or altering the rank order of pixel intensities. A white margin borders each inset, separating it from the underlying atlas lines. Numbers at the top left (01–16 in **a**, 01–12 in **b**) give the order along each series. **a**, Lateral (01, 02) to medial (15, 16), one lateral-to-medial step per row: the contralateral section on the left, its ipsilateral partner on the right; rostral is left and dorsal is up. **b**, Dorsal (01) to ventral (12); rostral is up and the hemispheres contralateral (Contra) and ipsilateral (Ipsi) to the inoculation are on the left and right of every section. The coronal plane is shown in Fig. 2b, Fig. 3 and Supplementary Fig. 3. Scale bars, 2 mm; each applies to all maps of its panel, not to the insets.

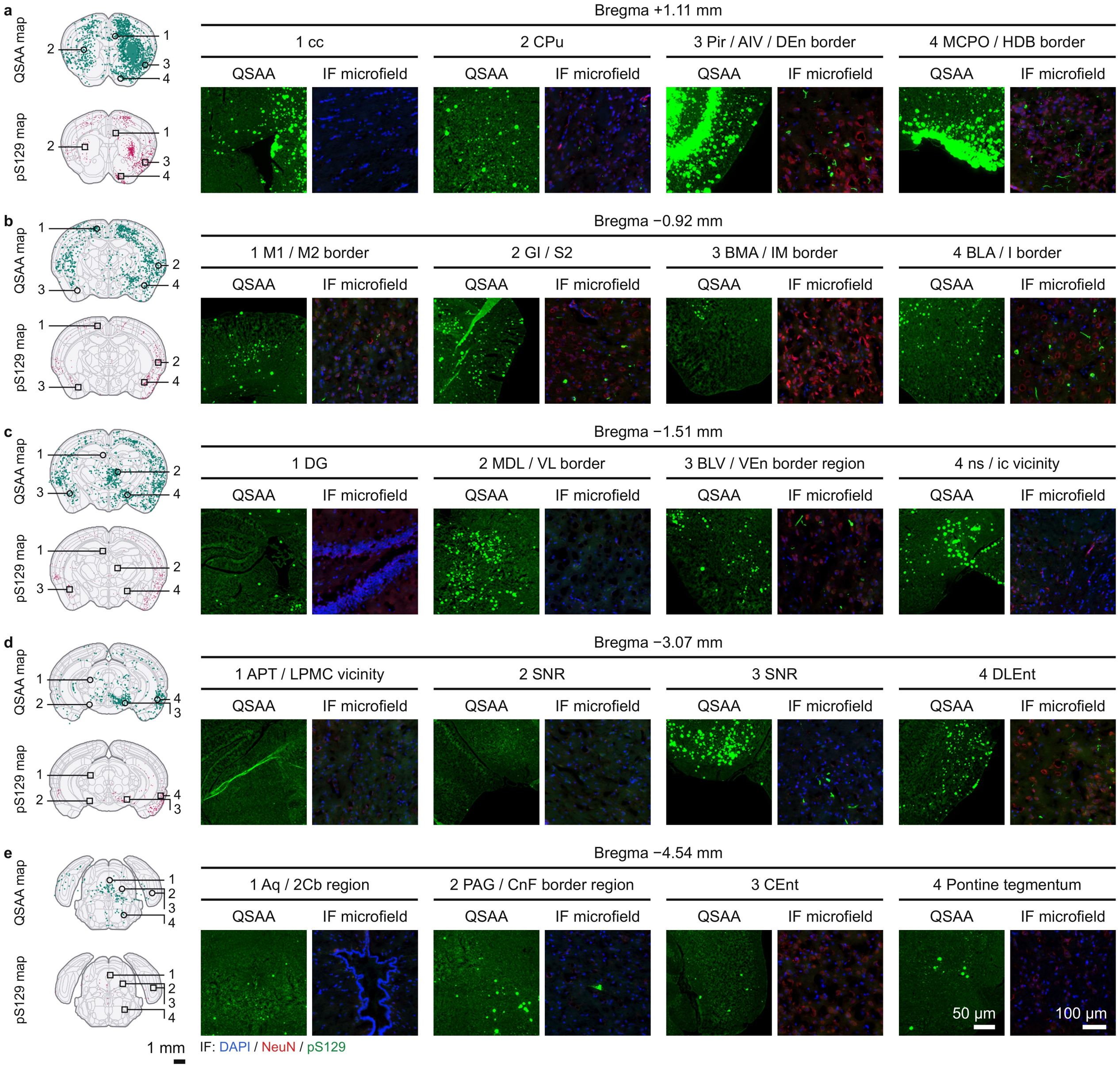


**Supplementary Fig. 3 | Object maps and enlarged fields compare QSAA products with pS129 immunofluorescence at five coronal levels of the mPFF-inoculated mouse brain.** **a**–**e**, Five coronal levels, rostral (**a**) to caudal (**e**), of mouse brain 180 days after inoculation of mouse α-synuclein (αSyn) preformed fibrils (mPFF) into one striatum. Headers, Bregma coordinate. Left column: QSAA map, the objects of the quiescent seed amplification assay (QSAA) on that level as teal dots on the atlas map; below, pS129 map, the objects of the matched section immunostained for αSyn phosphorylated at serine 129 (pS129) as magenta dots. Rings (QSAA map) and squares (pS129 map) mark the four fields enlarged on the right, each numbered beside the brain on a leader to its mark. A field is set at the position it has within the outline of its section, carried over to the outline of the atlas plate. Contralateral and ipsilateral hemispheres to the inoculation are on the left and right of every map. These maps come from the whole sections in Fig. 2b and Fig. 3a. Right: fields 1–4, named by region: QSAA, thioflavin T fluorescence of the QSAA products; IF microfield, the same region of the immunofluorescence section (pS129, green; neuronal nuclei antigen (NeuN), red; 4′,6-diamidino-2-phenylindole (DAPI), blue). Region names follow the Franklin and Paxinos atlas and are given in full in Supplementary Table 2. Scale bars, 1 mm (object maps), 50 µm (QSAA fields) and 100 µm (IF microfields).

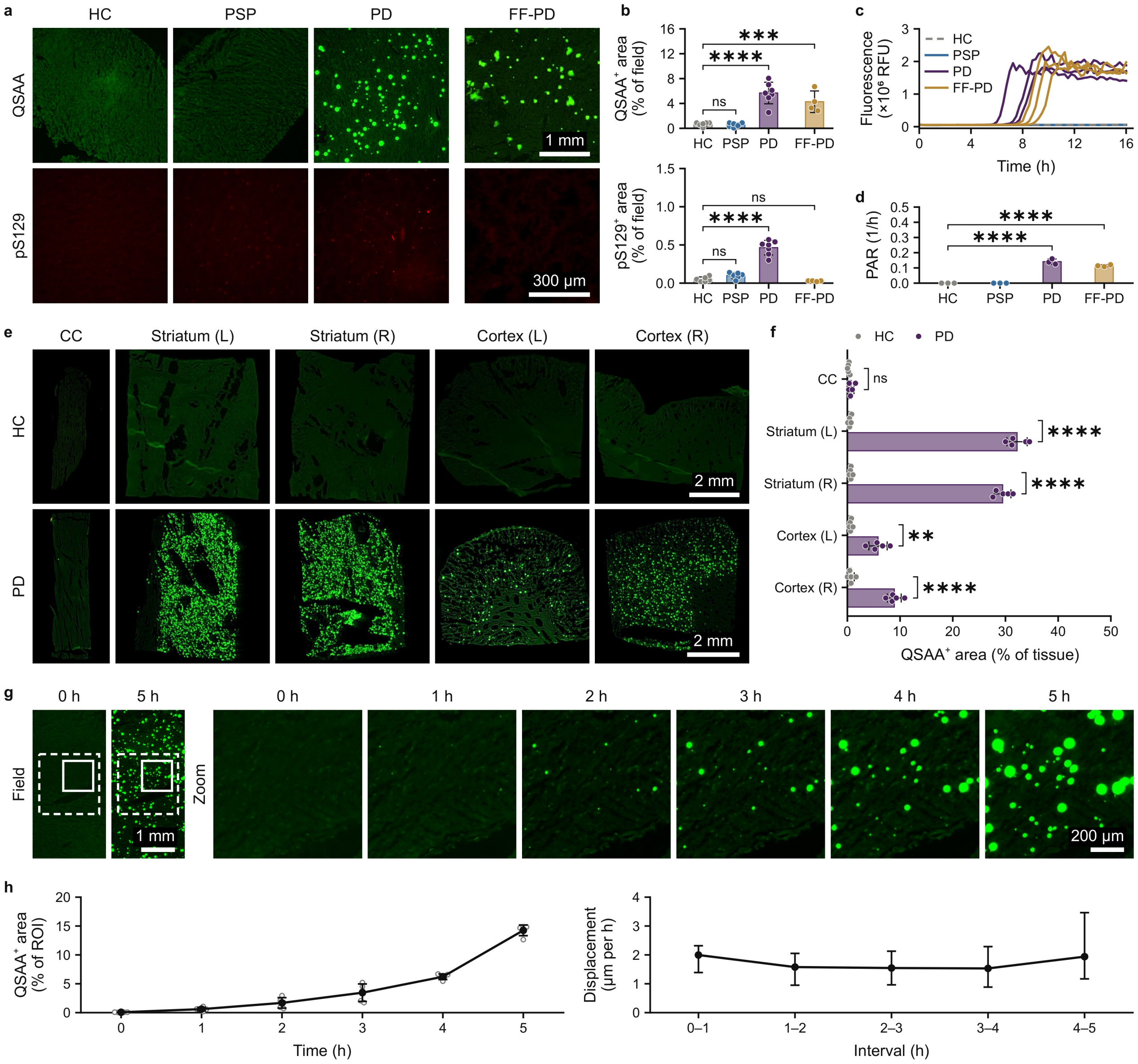


**Supplementary Fig. 4 | QSAA detects endogenous α-synuclein seeding activity in human Parkinson's disease brain sections, where the products grow in place.** **a**, Brain sections from a neurologically normal control (HC), a progressive supranuclear palsy case (PSP), a Parkinson's disease case (PD) and a long-term formalin-fixed PD specimen (FF-PD). Top row, thioflavin T (ThT) fluorescence after the quiescent seed amplification assay (QSAA; green); bottom row, immunofluorescence for α-synuclein phosphorylated at serine 129 (pS129; red). **b**, QSAA-positive (QSAA$^+$) and pS129-positive (pS129$^+$) area. Bars, means; dots, individual values; error bars, s.d.; for both QSAA$^+$ and pS129$^+$: PD n = 7, FF-PD n = 4 (technical replicates), PSP n = 6 and HC n = 6; one-way ANOVA with Dunnett's test against HC. **c**, Real-time quaking-induced conversion of brain lysates from the four groups; ThT fluorescence in relative fluorescence units (RFU). Each line, one reaction. **d**, Protein aggregation rate (PAR). Bars, means; dots, single reactions; error bars, s.d.; n = 3 per group; one-way ANOVA with Dunnett's test against HC. **e**, QSAA on sections from five sites of HC and PD brain: corpus callosum (CC), two striatal and two cortical sites. **f**, QSAA$^+$ area at the five sites. Bars, means; dots, individual values; error bars, s.d.; n = 5 per site and group; two-tailed unpaired *t*-test with Welch's correction, Holm-adjusted over the five sites. **g**, One field of a PD section during the reaction. Left (Field), 0 and 5 h; right (Zoom), solid box at 0–5 h; dashed box, region of interest (ROI) in **h**. **h**, ROI measurements in **g**. Left, QSAA$^+$ area (open circles, individual values; dots, means; error bars, s.d.; n = 5 per time point). Right, displacement of matched objects between consecutive images (dots, medians; error bars, interquartile ranges; n = 6, 29, 70, 98 and 63 objects for the five intervals). *$P < 0.05$, **$P < 0.01$, ***$P < 0.001$, ****$P < 0.0001$; ns, not significant. Scale bars, 1 mm (**a**, QSAA; **g**, Field), 300 μm (**a**, pS129), 2 mm (**e**, right cortex) and 200 μm (**g**, Zoom).

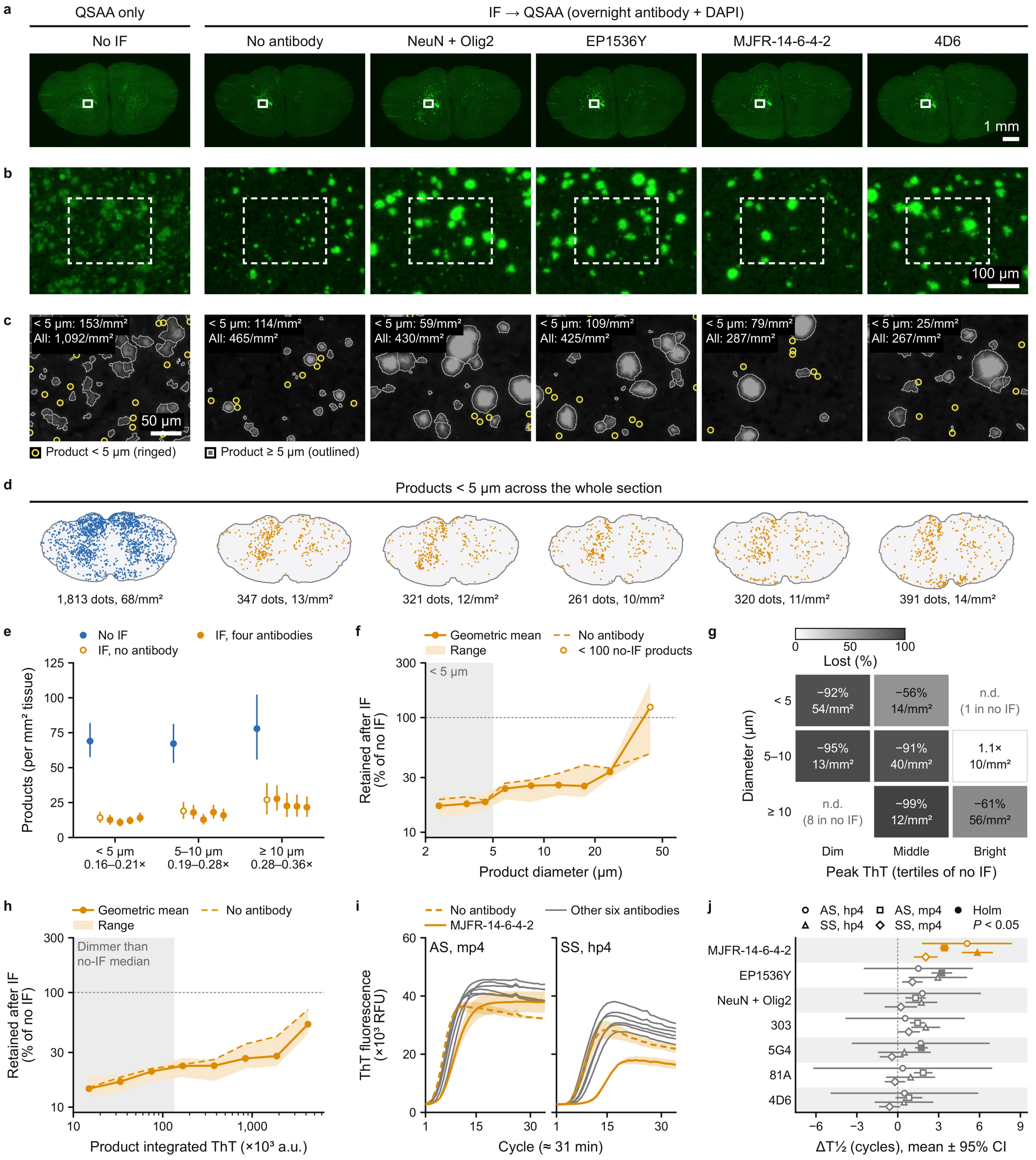


**Supplementary Fig. 5 | Immunofluorescence performed before QSAA removes most products, especially the small and dim ones, with or without antibody.** **a**, ThT fluorescence (green) of brain sections from mPFF-inoculated mice, after QSAA alone or preceded by IF without antibody or with the antibodies named (NeuN + Olig2, together); white boxes, fields in **b**. **b**, Dashed boxes, regions in **c**. **c**, ThT fluorescence (grey); yellow rings, products below 5 µm; outlines, larger products; tags, densities in **b**. **d**, One dot per product below 5 µm; blue, without IF; orange, after IF. **e**, Product density by size class; below each class, ratios to no IF. Circles, mean; error bars, 95% bootstrap confidence interval (CI); n = 92–101 tiles (500 µm) per section. **f**, Density after IF relative to no IF, by diameter: geometric mean and range of five IF sections. **g**, Density change after IF (upper value) by diameter and peak ThT intensity; lower value, density without IF; n.d., not determined. **h**, As **f**, by integrated ThT fluorescence. **i**, Seed amplification assay with antibody, in ammonium sulfate (AS) or sodium sulfate (SS). Lines, mean; bands, s.d. **j**, Time to half-maximal ThT fluorescence relative to no antibody (Δ T1/2). Symbols, difference of means; lines, 95% CI; filled, Holm-adjusted $P < 0.05$. n = 3 wells per group (**i**, **j**); two-tailed unpaired *t*-test with Welch's correction (**j**). Scale bars, 1 mm (**a**), 100 µm (**b**) and 50 µm (**c**).

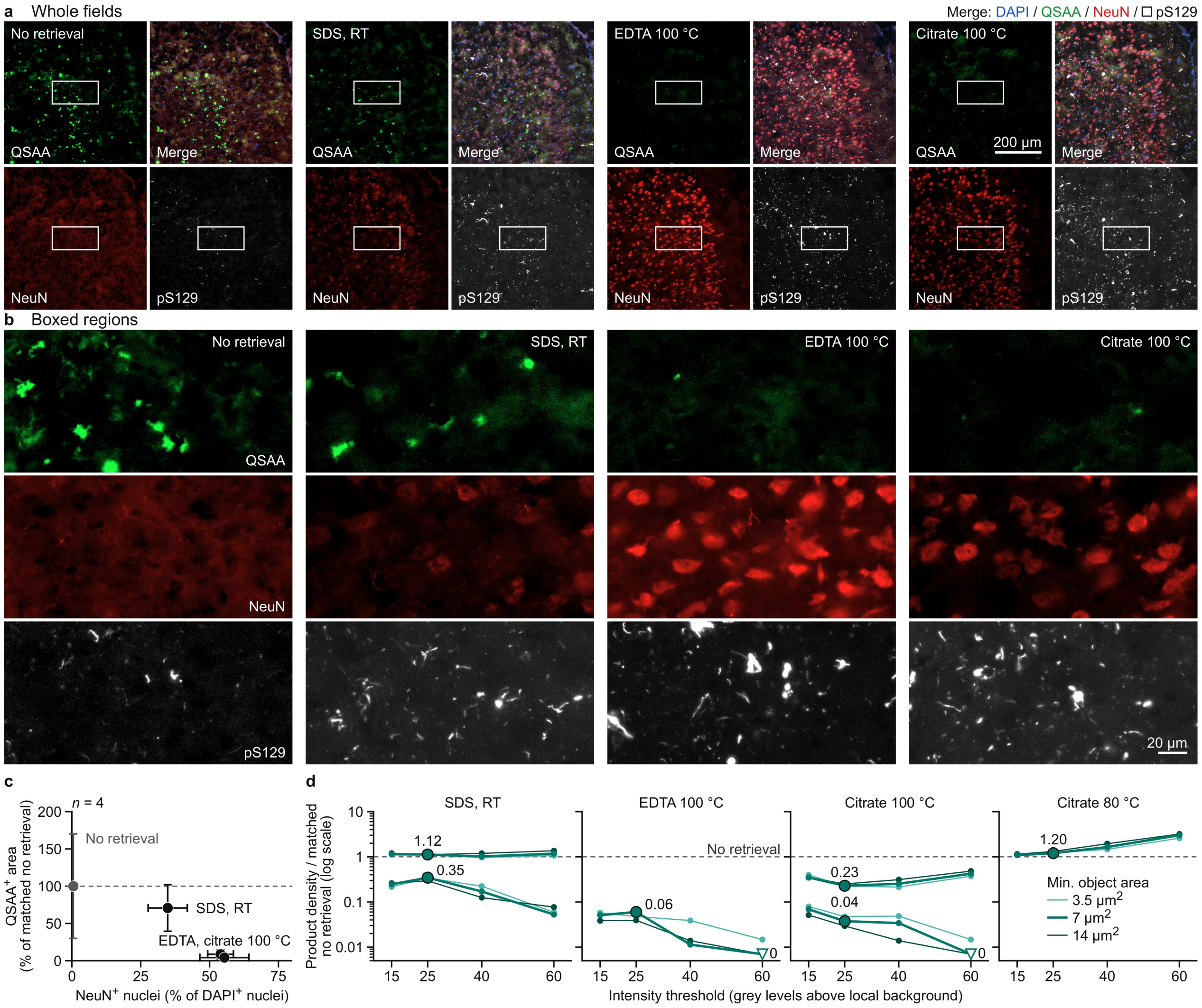


**Supplementary Fig. 6 | Antigen retrieval at 100 °C restores NeuN and pS129 immunofluorescence but removes most QSAA products.** **a**, Brain sections processed for the quiescent seed amplification assay (QSAA), then given no antigen retrieval, a sodium dodecyl sulfate (SDS)-based solution at room temperature (RT), or ethylenediaminetetraacetic acid (EDTA; pH 8.0) or citrate at 100 °C for 30 min, and immunostained for neuronal nuclei antigen (NeuN) and α-synuclein phosphorylated at serine 129 (pS129). One field per protocol: QSAA (thioflavin T fluorescence; green), NeuN (red), pS129 (grey scale) and the merge, with 4′,6-diamidino-2-phenylindole (DAPI) blue and pS129 white (open square in the key). **b**, White-boxed regions in **a**. **c**, QSAA-positive (QSAA$^+$) area against NeuN$^+$ nuclei (% of DAPI$^+$ nuclei) for the protocols of **a**. Grey, no retrieval; black, retrieval protocols. **d**, Density of QSAA products at four intensity thresholds (x axis) and three minimum object areas (lines; key). Citrate at 80 °C was also applied for 30 min. Large labelled circles, ratio at threshold 25 and minimum area 7 μm$^2$; open triangles, ratio 0. Matched no retrieval (**c**, **d**; dashed lines), no-retrieval section processed in parallel. Data are mean ± s.d.; n = 4 independent replicates per protocol. Scale bars, 200 μm (**a**) and 20 μm (**b**).

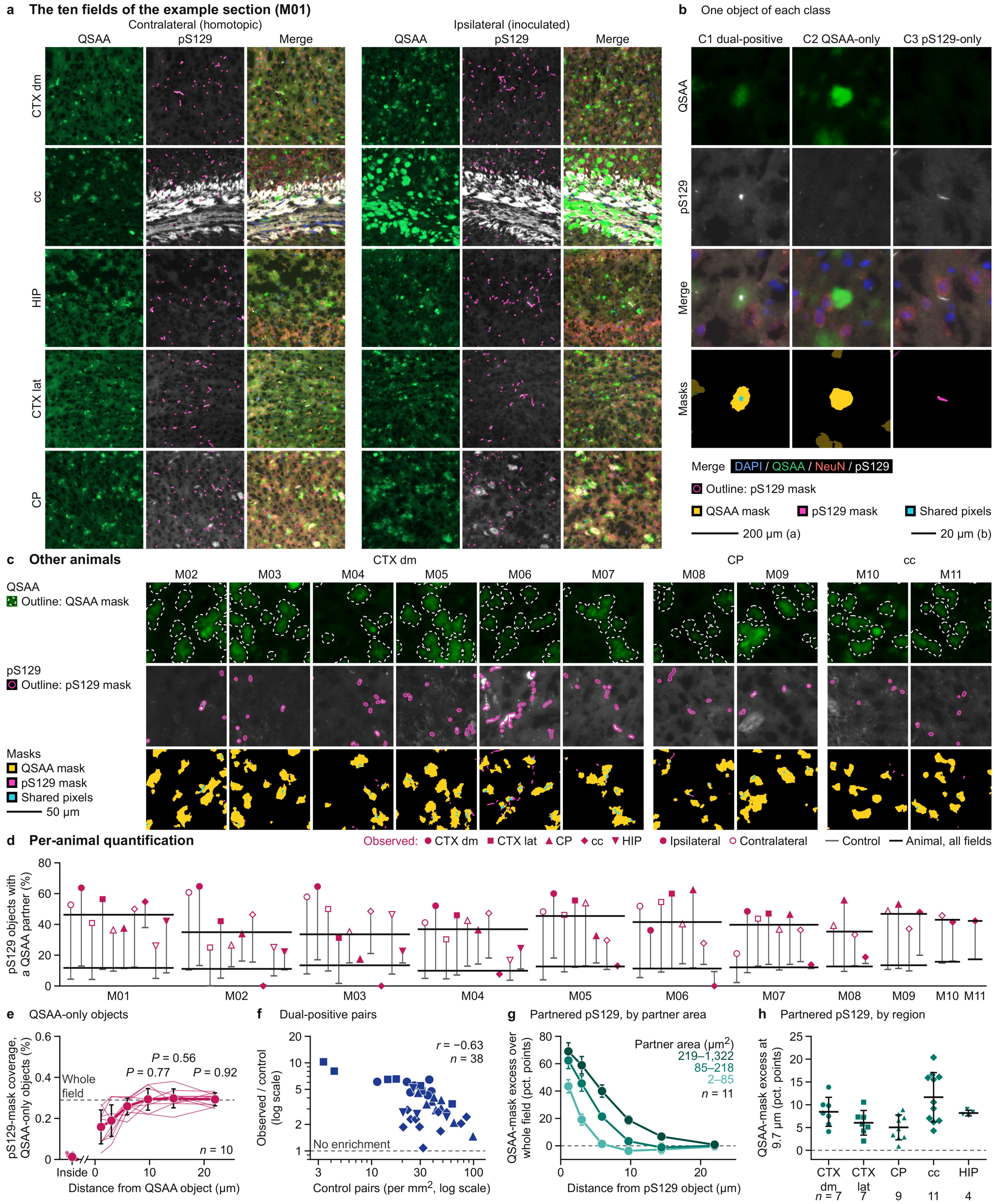


**Supplementary Fig. 7 | Analysis of QSAA–pS129 object pairing across fields, animals and brain regions.** **a**, Ten fields of the Fig. 7a section (M01), contralateral (Contra) and ipsilateral (Ipsi): QSAA, pS129 and merge (4′,6-diamidino-2-phenylindole (DAPI), blue; QSAA, green; neuronal nuclei antigen (NeuN), red; pS129, white). Rows, CTX dm, cc, HIP, CTX lat and CP. Magenta outlines, pS129 objects counted. **b**, One object per Fig. 7c class; masks: QSAA, yellow; pS129, magenta; shared pixels, cyan; others dimmed. **c**, One ipsilateral field per other animal (M02–M11), as in **b**; dashed black-and-white outlines, QSAA mask, 3.8 µm outside it. **d**, pS129 objects with a QSAA partner in the 75 fields (Fig. 7d): symbols, observed (shape, region; filled, Ipsi; open, Contra); grey ticks, control; black lines, each animal's values over its fields. **e**, pS129-mask coverage inside and around QSAA-only objects; thin lines, animals; P = 0.77, 0.56 and 0.92. **f**, Observed/control ratio of dual-positive pairs against control pair density (n = 38 animal–region values); r, Pearson correlation of logs. **g**, QSAA-mask excess around partnered pS129 objects, by thirds of QSAA-partner area. **h**, The same at 9.7 µm, by region; symbols, animals. Data in **e**, **g** and **h**, mean ± s.d.; n = 11 animals; exact two-sided Wilcoxon signed-rank test against each animal's whole-field fraction (**e**). Scale bars, 200 µm (**a**), 20 µm (**b**) and 50 µm (**c**).

### Supplementary Tables

**Supplementary Table 1 | Key reagents and resources.**
Antibodies, chemicals and buffers, consumables and columns, instruments, and software and algorithms used in this study.

| Reagent or resource | Source | Identifier / catalogue no. | Application / dilution |
|---|---|---|---|
| **Antibodies** | | | |
| Rabbit monoclonal anti-pS129-αSyn (clone EP1536Y) | Abcam | ab51253 | IF (1:1,000); SAA (1:100) |
| Mouse monoclonal anti-pS129-αSyn (clone 81A) | Abcam | ab184674 | IF (1:1,000) |
| Rabbit monoclonal anti-αSyn filament (clone MJFR-14-6-4-2) | Abcam | ab209538 | IF / SAA (1:100–1:1,000) |
| Mouse monoclonal anti-αSyn (clone 4D6) | Abcam | ab1903 | IF / SAA (1:100–1:1,000) |
| Mouse monoclonal anti-NeuN (clone A60) | Millipore | MAB377 | IF (1:1,000); SAA (1:100) |
| Rabbit polyclonal anti-IBA1 | FUJIFILM Wako | 019-19741 | IF (1:1,000) |
| Mouse monoclonal anti-GFAP | Cell Signaling Technology | 3670 | IF (1:1,000) |
| Goat anti-Rabbit IgG (H+L), Alexa Fluor 488 | Invitrogen | A-11008 | Standalone IF (1:1,000) |
| Goat anti-Mouse IgG (H+L), Alexa Fluor 488 | Invitrogen | A-11001 | Standalone IF (1:1,000) |
| Goat anti-Rabbit IgG (H+L), Alexa Fluor 555 | Invitrogen | A-21428 | Multiplex IF-QSAA (1:1,000) |
| Goat anti-Mouse IgG (H+L), Alexa Fluor 555 | Invitrogen | A-21422 | Multiplex IF-QSAA (1:1,000) |
| Goat anti-Rabbit IgG (H+L), Alexa Fluor 647 | Invitrogen | A-21245 | Multiplex IF-QSAA (1:1,000) |
| Goat anti-Mouse IgG (H+L), Alexa Fluor 647 | Invitrogen | A-21235 | Multiplex IF-QSAA (1:1,000) |
| **Chemicals, buffers and reagents** | | | |
| Thioflavin T (ThT) | Sigma-Aldrich | T3516 | 40 μM (SAA / QSAA) |
| Thioflavin S (ThS) | Sigma-Aldrich | T1892 | 0.05% (amyloid staining) |
| Sodium sulfate ($Na_2SO_4$) | Sigma-Aldrich | 238597 | 800 mM (QSAA buffer) |
| Ammonium sulfate ($(NH_4)_2SO_4$) | Sigma-Aldrich | A4915 | Protein purification / SAA |
| Agarose | BioFroxx | 1110GR100 | 1% (w/v) tissue hydrogel overlay |
| SDS-based antigen retrieval solution | Beyotime | P0090 | Antigen retrieval comparison |
| TrueBlack Lipofuscin Autofluorescence Quencher | Biotium | 23007 | Autofluorescence quenching |
| Mineral oil | Biosharp | BS927 | Quiescent plate-based SAA overlay |
| Protease inhibitor cocktail | Roche | 04693116001 | Recombinant protein lysis |
| High-capacity endotoxin removal resin | Sangon Biotech | B641718 | Endotoxin removal |

| Reagent or resource | Source | Identifier / catalogue no. | Application / dilution |
| --- | --- | --- | --- |
| **Consumables and columns** | | | |
| High-adhesion glass slides | CITOTEST | 188105W | Cryosection mounting |
| 96-well black, clear-bottom microplate | Nunc / Thermo Scientific | 165305 | Agitated SAA |
| 96-well polypropylene PCR plate | Thermo Scientific | AB0700 | Quiescent SAA |
| Optical adhesive sealing film | Applied Biosystems | 4311971 | Plate sealing |
| HiTrap Q HP anion-exchange column | Cytiva | 17115401 | αSyn monomer purification |
| Amicon Ultra 50-kDa MWCO filter | Millipore | UFC905096 | Monomer ultrafiltration |
| 10-µL syringe with 33-gauge needle | Hamilton | 65460-05 | Stereotaxic injection |
| **Instruments and equipment** | | | |
| Stereotaxic instrument | Kopf Instruments | Model 940 | Mouse stereotaxic surgery |
| ThermoMixer C | Eppendorf | 5382 | Fibril generation |
| Non-contact water-bath sonicator | SCIENTZ | SCIENTZ08-III | PFF fragmentation |
| Precision temperature-controlled stage | Ruicheng Instrument Co. | TS400 | In situ slide QSAA |
| Multimode microplate reader | Molecular Devices | SpectraMax iD5 | Plate-based SAA kinetics |
| Real-time fluorescence thermal cycler | Bio-Rad | CFX96 | Quiescent plate-based SAA |
| Fluorescence microplate reader | BMG LABTECH | FLUOstar Omega | Brain homogenate RT-QuIC |
| Inverted widefield fluorescence microscope | Nikon | Eclipse Ti2-E (Digital Sight 50M) | Whole-section and field imaging |
| Transmission electron microscope | Thermo Fisher / FEI | Talos L120C | Negative-stain and in situ TEM |
| **Software and algorithms** | | | |
| NIS-Elements Advanced Research | Nikon | v5.30 | Widefield image acquisition |
| Fiji / ImageJ | NIH | v2.3.0 | Image inspection and preprocessing |
| Custom Python spatial analysis pipeline | This study | FluoroAnts v6.3 / OpenCV / scikit-image / SciPy | Unmixing, segmentation, atlas mapping & null models |
| GraphPad Prism | GraphPad Software | v10.2.2 | Statistical analysis |

**Supplementary Table 2 | Region and tract abbreviations.**

Abbreviations used in Fig. 3, Fig. 7, Supplementary Fig. 3 and Supplementary Fig. 7. Region names follow the Franklin and Paxinos mouse brain atlas or the Allen reference atlas.

| Abbreviation | Full name |
|---|---|
| 2Cb | second cerebellar lobule |
| ACB | nucleus accumbens |
| AId | dorsal agranular insular area |
| AIV | ventral agranular insular cortex |
| alv | alveus |
| APT | anterior pretectal nucleus |
| Aq | aqueduct |
| BLA | anterior basolateral amygdaloid nucleus |
| BLV | ventral basolateral amygdaloid nucleus |
| BMA | anterior basomedial amygdaloid nucleus |
| CA1 | hippocampal field CA1 |
| CA3 | hippocampal field CA3 |
| cc | corpus callosum |
| ccb | body of the corpus callosum |
| ccg | genu of the corpus callosum |
| CEnt | caudomedial entorhinal cortex |
| cing | cingulum bundle |
| CnF | cuneiform nucleus |
| Contra | contralateral to the inoculation |
| CP | caudoputamen |
| CPu | caudate putamen |
| CTX dm | dorsomedial isocortex |
| CTX lat | lateral isocortex |
| DEn | dorsal endopiriform nucleus |
| DG | dentate gyrus |
| DI | dysgranular insular cortex |
| DLEnt | dorsolateral entorhinal cortex |
| Dorsal wm band | dorsal white-matter band, combining cc *sensu lato* (fa, ccg and ccb), cing and scwm |
| ec | external capsule |
| ENTl | lateral entorhinal area |
| fa | anterior forceps of the corpus callosum |
| gcc | genu of the corpus callosum, as labelled in Fig. 3b |
| GI | granular insular cortex |
| HDB | horizontal limb of the diagonal band |
| HIP | hippocampus |
| I | intercalated amygdaloid nuclei |

| Abbreviation | Full name |
|---|---|
| ic | internal capsule |
| IM | main intercalated amygdaloid nucleus |
| Ipsi | ipsilateral to the inoculation |
| LPMC | mediocaudal lateral posterior thalamic nucleus |
| M1 | primary motor cortex |
| M2 | secondary motor cortex |
| MCPO | magnocellular preoptic nucleus |
| MDL | lateral mediodorsal thalamic nucleus |
| ml | medial lemniscus |
| MOp | primary motor area |
| MOs | secondary motor area |
| ns | nigrostriatal bundle |
| opt | optic tract |
| OT | olfactory tubercle |
| PAG | periaqueductal grey |
| PIR | piriform area |
| Pir | piriform cortex |
| RSPv | ventral retrosplenial area |
| S2 | secondary somatosensory cortex |
| scwm | supra-callosal cerebral white matter |
| SNR | reticular part of the substantia nigra |
| SSp | primary somatosensory area |
| SSs | supplemental somatosensory area |
| VEn | ventral endopiriform nucleus |
| VISC | visceral area |
| VISp | primary visual area |
| VL | ventrolateral thalamic nucleus |
